# The long non-coding RNA ACHLYS modulates biomolecular condensates to regulate alternative splicing in root development

**DOI:** 10.64898/2026.02.20.706954

**Authors:** M. Heidecker, P. Mammi, T. Bergelt, A. Christ, M. Lewinski, T. Köster, C. Charon, Y. Jin, M. Marquardt, T. Blein, J. Bazin, D. Staiger, M. Crespi

## Abstract

Alternative splicing (AS) enables eukaryotes to dynamically adjust RNA and protein isoforms encoded in one gene. Long non-coding RNAs (lncRNA) have emerged as novel regulators of AS through multiple modes of action, including interactions with splicing factor (SF) proteins. Here, we used *Arabidopsis thaliana* lateral root development to dissect the specificity of lncRNAs in AS regulation. Data mining and a transient expression screen allowed us to identify novel lncRNAs interacting *in vivo* with the well-characterized SFs Nuclear speckle RNA binding protein A (NSRa) and glycine-rich RNA-binding protein 7 (GRP7), that are differentially expressed during lateral root organogenesis. We identified 4 lncRNAs affecting more than 250 AS events in plant cells, whereby most AS events are unique for each lncRNA. Notably, 19% of the AS events linked to two NSRa-recognized lncRNAs are identical and positively correlated. One lncRNA was named *ACHLYS* and is highly conserved in several *Brassicaceae* whereas another was the recently characterized *FLAIL* lncRNA. *ACHLYS* knockdown (KD-*ACHLYS*) and overexpressing lines (OE-*ACHLYS*) modulate AS regulation in plants and affect root architecture. In parallel, we performed NSRa-iCLIP to determine *in vivo* genome-wide NSRa binding sites. Interestingly, NSRa binding sites are locally enriched in *ACHLYS*-dependent AS introns and *ACHLYS* itself. A specific *ACHLYS*-induced AS event requires NSRa function showing that *ACHLYS* defines AS targets through NSRa interaction on specific transcripts. *In vitro* assays indicate that direct interaction of NSRa with *ACHLYS* promotes NSRa phase separation. Furthermore, overexpression of *ACHLYS* in NSRa-GFP plants results in increased accumulation of NSRa in Nuclear speckles. Our data revealed that *ACHLYS* can modulate NSRa condensates *in vivo* and binds to NSRa for modifying AS patterns, suggesting a new mechanism where lncRNAs can fine tune SFs regulation and the transcriptome output in eukaryotic organogenesis.

## Introduction

Splicing is the central processing step of precursor-(pre-) mRNA in all eukaryotes and allows the correct expression of genetic information as mature RNA. The spliceosome, a dynamic ribonucleoprotein complex, assembles at the intron/exon junctions known as 5’- and 3’-splice sites (SS) and catalyzes the intron removal and ligation of the flanking exons (Plaschka et al. 2019). Splicing is highly controlled on multiple levels and depending on regulatory factors, different SSs in the same pre-mRNA may or may not be selected by the spliceosome, resulting in different mature RNAs. This process is called alternative splicing (AS). In humans ∼90%, and in the model plant *Arabidopsis thaliana,* ∼60% of all intron-containing genes undergo AS, resulting in an increased transcriptome and proteome complexity (Chaudhary et al. 2019). Thereby, AS contributes to tissue identity, organ development and environmental responses (Syed et al. 2012; Staiger and Brown 2013; Alhabsi et al. 2025). Key AS regulators are RNA binding proteins, called splicing factors (SFs). The two major types of SFs are serine/arginine-rich proteins (SRs) and heterogeneous nuclear ribonucleoproteins (hnRNPs). Multiple SFs bind the pre-mRNA to repress or promote spliceosome assembly and thereby affect SS usage (Ule and Blencowe 2019). The modulation of SF expression, subcellular localization and posttranslational modifications is an integral part of AS regulation (Stamm 2008).

Long non-coding RNAs (lncRNAs) are transcripts longer than 200 nt with no or low coding potential. Biological functions of lncRNAs are described through plant development and as response to biotic and abiotic stimuli. Their main molecular function is to fine tune gene expression regulation by direct interaction with proteins, for instance chromatin remodelers, transcription factors and RNA binding proteins involved in RNA processing steps like AS (Ariel et al. 2015; Statello et al. 2021). lncRNAs regulate AS through different mechanisms, often by interacting with SFs. For example, lncRNAs can block, decoy and recruit SFs, or modulate SF activity (Romero-Barrios et al. 2018).

Two plant specific SFs are Nuclear speckle RNA binding protein a (NSRa) and glycine-rich RNA-binding protein 7 (GRP7). NSRa and its homologue NSRb are SR-like SFs (Lucero et al. 2020). NSRa was found to bind hundreds of transcripts and to regulate AS (Bazin et al. 2018). Notably, NSRa co-localizes with other SFs to Nuclear speckles, a biomolecular condensate involved in AS regulation (Bardou et al., 2014). GRP7 and its homologue GRP8 are well-characterized as hnRNP-like SFs. It was shown that GRP7 binds hundreds of mRNAs, preferentially at the 3’ untranslated regions (3’-UTRs) and regulates AS, stability, and other RNA processing steps (Streitner et al. 2012; Koester et al. 2014; Meyer et al. 2017; Lewinski et al. 2024). The RNA interactome of NSRa and GRP7 showed that the two SFs not only bind to mRNA but also to hundreds of lncRNAs. Nevertheless, the functional role of this interaction remains unexplored (Meyer et al. 2017; Bazin et al. 2018). However, most studies on lncRNA-mediated regulation of AS have been performed in cell culture, particularly in cancer cells, and very little is known about their roles in complex organisms under physiological conditions.

AS contributes to the definition of tissue identity in both humans and plants (Baralle and Giudice 2017; Martín et al. 2021; Mazin et al. 2021). In plants, one central organ is the root, which is continuously exposed to a complex and changing environment. A dynamic response of the root system architecture to the environment is crucial for plant survival and is mainly defined by the number and length of lateral roots. Lateral root formation (LRF) is induced at specific xylem pole pericycle and controlled cell division process lead to the formation of a lassteral root meristem which physically penetrates the main root endodermis, cortex and epidermis to further develop as an organ and fulfill its biological function (Goh et al. 2023). LRF has been shown to be modulated by specific AS events (Remy et al. 2013; Jayaweera et al. 2014). A global role of AS in root development has been proposed by cell type specific transcriptome and proteome analysis (P. Lan et al. 2013; Li et al. 2016).

Two NSRa-bound lncRNAs, *ALTERNATIVE SPLICING COMPETITOR* (*ASCO*) and *FLAIL*, were characterized as AS regulators in *Arabidopsis thaliana* (Bardou et al. 2014; Rigo et al. 2020; Jin et al. 2023). Nevertheless, the mechanism by which lncRNAs interaction with SFs triggers specific AS changes is not well characterized, raising the question whether lncRNAs are a general component of AS networks and their specificity.

In this work, we combined a high-resolution time-course transcriptome analysis of LRF with *in vivo* binding site determination of SFs to identify lncRNA candidates that may be involved in AS regulation during LRF. The effect of lncRNA candidates on AS was tested in a transient expression assay and led to the identification of a new lncRNA, we named *ACHLYS*. *ACHLYS* knockdown or overexpression impairs root development and affects AS. In parallel, NSRa individual nucleotide-resolution crosslinking and immunoprecipitation (iCLIP) allowed us to identify NSRa binding sites on *ACHLYS* and to show that NSRa binding sites are enriched in *ACHLYS*-dependent AS events. We further demonstrate that individual AS events induced by transient *ACHLYS* expression in wild type plants are absent in the *nsra* mutant and that NSRa containing biomolecular condensates are regulated by *ACHLYS*. This data suggest that *ACHLYS* modulates AS through interaction with NSRa and that lncRNAs directly interacting with SFs are general components of dynamic AS regulation in plants.

## Results

### Alternative splicing landscapes across lateral root formation

The formation of a lateral root can be induced by a gravitropic stimulus after 90° rotation of *A. thaliana* seedlings. The lateral root develops in a synchronized manner and the hours after gravitropic stimulus correspond to different developmental stages of the lateral root primordia (Lucas et al. 2008). To address the contribution of AS in lateral root organogenesis, we exploited RNAseq data of LRF, induced by gravitropic stimulus, spanning nine time points every 6 hours from T0 to T48 (Figure 1A). To identify individual splicing events during LRF we used SUPPA2 (Trincado et al. 2018) and compared the splicing profile of each time point to T0. We detected a total of 3113 differential AS (DAS) events. Already in the first 24 h of LRF we detected 200 AS events and from T36-T48 more than 500, the majority are alternative 3’-SS (35%) 5’-SS (13-17%) and intron retention (23-35%) (Figure 1B).

**Figure 1:**
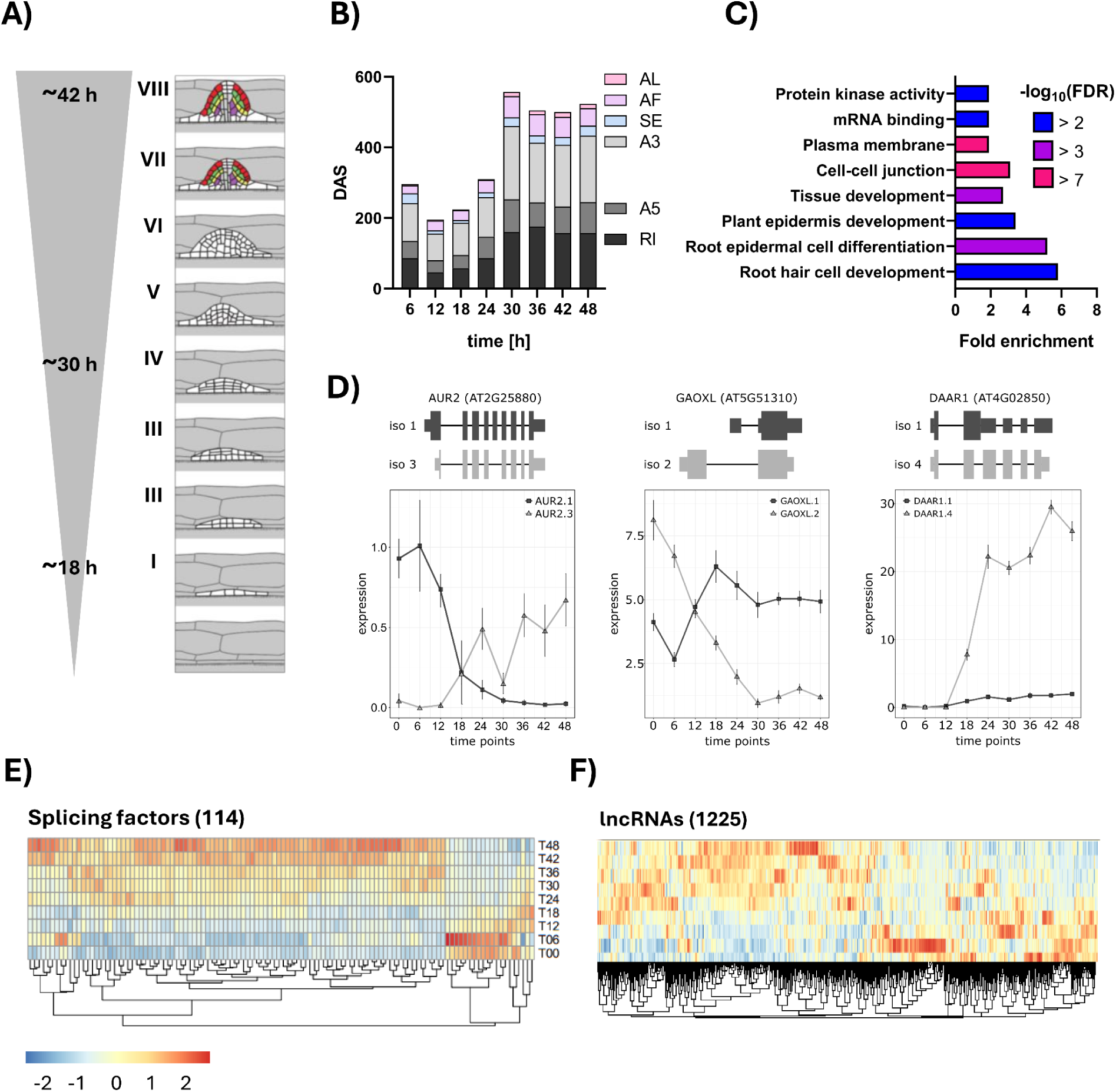
Alternative splicing events and gene expression patterns of splicing factors and lncRNAs during lateral root formation. **A)** Scheme of developmental stages of lateral root primordium formation up to 42 h after root bending. Image adapted from Péret et al. 2012. **B)** Number of differential alternative splicing (DAS) events during the time points depicted in 1A from lateral root formation (LRF) as determined by SUPPA2. Splicing events are classified as Retained introns (RI) alternative 3’ splice site (A3) alternative 5’ splice site (A5), alternative first exon (AF), alternative last exon (AL) and skipped exon (SE). **C)** Gene ontology analysis of all DAS genes detected during LRF. **D)** Schematic representation of alternative spliced isoforms. Bars represent Exons and lines, introns; narrow bars represent 5’- and 3’-UTRs, and wide bars represent CDS. Graphs show expression levels of individual splicing isoforms during LRF timepoints, indicating isoform switches for αAurora2 (AUR2), 2-oxoglutarate and Fe(II)-dependent oxygenase superfamily protein (GAOXL) and D-AMINO ACID RACEMASE1 (DAAR1). **E-F)** Hierarchical clustering of splicing regulators and lncRNAs along LRF. Heatmap of Z-score calculated based on log_2_ fold change determined by differential gene expression analysis.

Gene ontology (GO) analysis revealed that AS genes are enriched in GO terms related to root development, like “root hair cell development” (False Discovery Rate (FDR) = 1. 29*10^-3^), “root epidermal cell differentiation” (FDR = 3.98*10^-4^), “plant epidermis development” (FDR = 1.29*10^-3^) and “tissue development” (FDR = 1.7*10^-4^) (Figure 1C).

To identify switches of mRNA isoforms that may lead to proteins with different domain composition, throughout the entire time course of LRF we used TSIS (J.-C. Guo et al. 2019) and identified 202 isoform switches in 119 genes. Isoform switching was most prominent during the mid-initiation stage (T18–T30), with a peak around 20 hours. These AS events were found in genes with diverse predicted functions, including 8 transcription factors, 27 RNA-binding proteins, and 18 genes linked to the GO term “root”. For instance, an alternative first exon and exon skipping of the α Aurora Kinase mRNA (AUR2, AT2G25880) results in decreased AUR2.1 expression, which encodes the full-length protein, and increased AUR2.3 expression, which lacks the Ser/Thr kinase active site of the phosphotransferase domain (Figure 1D). Double knockout mutants of AUR2 and its closest homologue AUR1 showed reduced lateral root density (Van Damme et al. 2011). Another example of a transcript associated with root development is the Fe(II)-dependent oxygenase superfamily protein GAOXL (AT5G51310) (Lin et al. 2011). The selection of an alternative first exon of GAOXL leads to increased GAOXL.1 and decreased GAOXL.2 isoform expression during LRF (Figure 1D).

To identify potential AS regulators during LRF, we searched for differentially expressed known AS regulators such as SFs and lncRNAs. We identified 114 SFs and 1225 lncRNAs that are differentially regulated during LRF and clustered them based on their expression pattern (Figure 1E,F). The majority of SFs were induced during late initiation and emergence stage (T18-T48). However, a small cluster showed an early-stage specific expression pattern with a peak at 6h. Notably, the SF NSRa and GRP7, for which the RNA interactome was identified, and their homologues NSRb and GRP8 are differentially expressed during LRF (Meyer et al. 2017; Bazin et al. 2018) (Figure S1A). They show different expression patterns while NSRa expression peaks at early and GRP7 at late time points. Globally, lncRNAs differentially expressed during LRF show more diverse time-point specific peaks than SFs, harboring the potential for contributing to point specific AS regulation (Figure 1E,F).

### Screening for lncRNAs involved in alternative splicing regulation identifies ACHLYS

To identify lncRNAs involved in AS regulation during LRF, we performed a bioinformatic screen for lncRNAs that interact with the plant specific SR-like SF NSRa and the hnRNP-like SF GRP7. First, the transcripts interacting with NSRa determined by RNA Immunoprecipitation and sequencing (RIPseq) (Bazin et al. 2018) and with GRP7 determined by individual-nucleotide resolution crosslinking and immunoprecipitation (iCLIP) (Meyer et al. 2017) were intersected with a manually curated list of *A. thaliana* lncRNAs. We identified 28 lncRNAs out of 342 NSRa-targets (8%) and 103 lncRNAs out of 858 GRP7-targets (12%). Subsequently, we filtered for lncRNAs without annotated antisense transcripts and introns in TAIR10 reference genome and manually excluded precursors of small RNAs to enrich for lncRNAs possible acting through AS regulation. Finally, we selected those lncRNAs that are differentially expressed in the LRF kinetics. In this way, we identified five lncRNAs that interact with NSRa (1- to 5-NSRa) and six lncRNAs that interact with GRP7 (1- to *6-GRP7*). As controls we selected two lncRNAs that were not recognized by the SFs NSRa and GRP7 (not bound by SFs (NBS) 1 and 2). For each of the 13 lncRNA candidates, characteristics including lengths, coding probability, expression levels in root, shoot and leaves, as well as expression during LRF, are summarized in Figure S1A.

To test the effect of the 13 lncRNA candidates on AS *in planta* we performed an *Agrobacterium*-mediated transient expression assay followed by RNAseq in *A. thaliana* leaves (Zhang et al. 2020). We overexpressed each lncRNA, *ASCO* as positive control and green fluorescent protein (GFP) as negative control, in the seventh leaf of *A. thaliana* plants. The NSRa-interacting lncRNAs *1-NSRa* and *2-NSRa*, *ASCO*, all six GRP7 interacting lncRNAs and control lncRNAs were successfully expressed (Figure S1B). Strand-specific libraries from poly(A)-RNA for four biological replicates were sequenced for these eleven lncRNAs and the GFP control.

To assess if transient expression of the lncRNAs has global effects on transcript levels we used DESeq2 to detect differentially expressed genes (DEGs). Interestingly, the effect of individual lncRNAs has a positive and negative effect on mRNA levels and ranked from less than 25 DEGs (5-GRP7 and *6-GRP7*), to over several hundred (*1-NSRa*, *1-NBS* and *2-NBS*) and up to thousands (*ASCO*, *2-NSRa*, *1-GRP7*, *2-GRP7* and *4-GRP7*) (Figure 2A).

**Figure 2:**
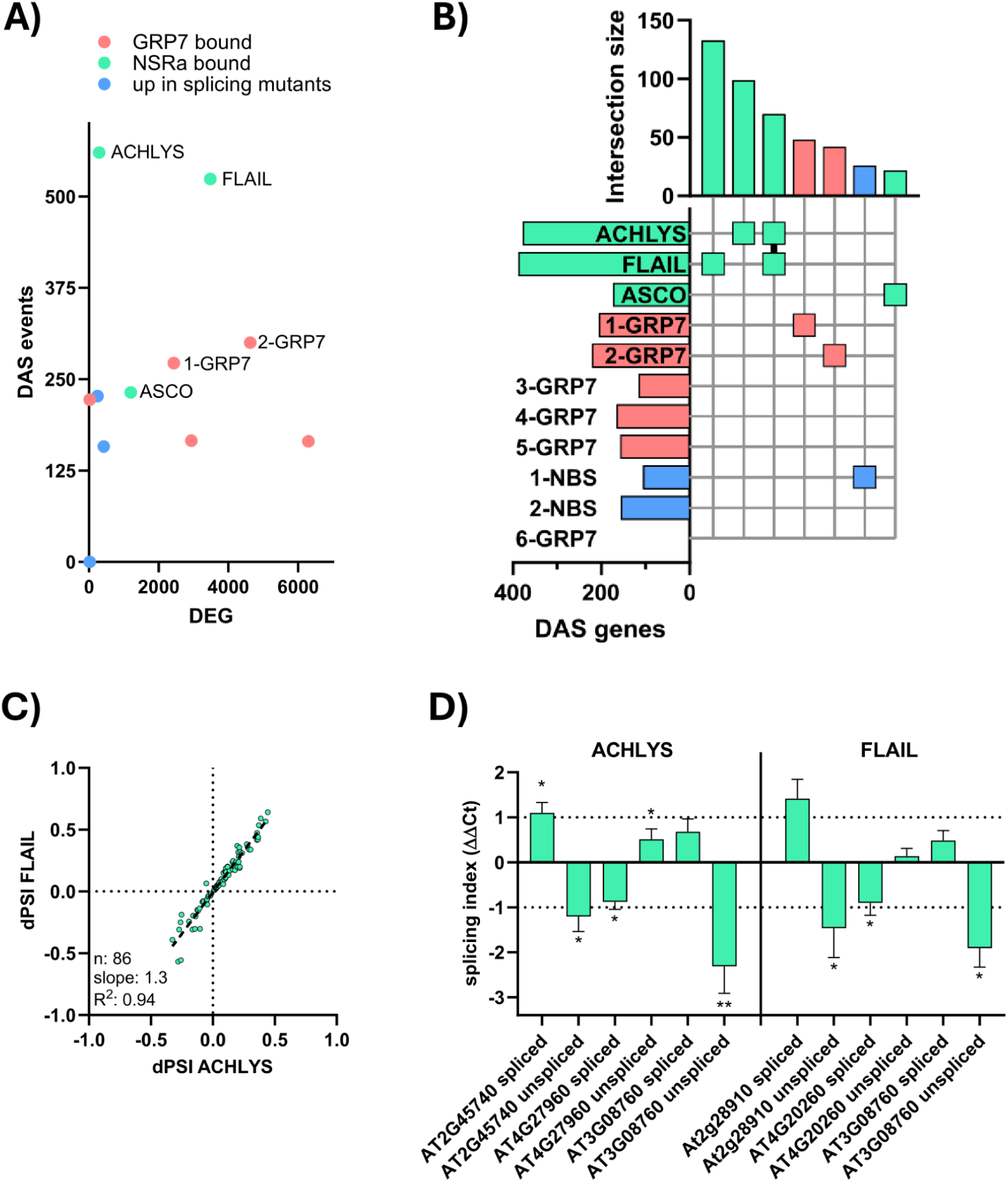
Identification of lncRNAs regulating splicing by a transient expression screen. **A)** Number differentially expressed genes (DEG) and differential alternative spliced (DAS) events after transient lncRNA expression compared to a GFP control. **B)** Upset graph showing first seven results after intersection of all DAS gene list for 11 transiently expressed lncRNAs, names are mentioned in results, red, green and blue color are those identified in GRP7 iCLIP data, NSRa RIPseq or control not bound by SFs (NBS) lncRNAs. **C)** Correlation between *ACHLYS* and *FLAIL* delta percent spliced in (dPSI or splicing index) of common DAS events identified in the intersection of Fig. 1B. Grey dots indicate splicing events confirmed by qPCR. Dotted black line displays linear regression model based on individual DAS events (dots). Number of DAS events (n), slope and R2 of the linear regression are shown. **D)** RNA levels of spliced and intron retention isoforms after transient expression of *ACHLYS* and *FLAIL* compared to GFP, quantified by qPCR. The splicing index ΔΔCt is the mean difference (ΔCt lncRNA minus ΔCt GFP comparing 3 to 4 biological replicates) and standard error of mean (unpaired, two-tailed T-test, *p ≤ 0.05, **p ≤ 0.01).

To quantify changes in AS pattern we performed Splicing profile comparison using transcript level quantification from the Arabidopsis Thaliana Reference Transcript Dataset 2 (AtRTD2) (R. Zhang et al. 2017) coupled with SUPPA2 (Trincado et al. 2018), which calculates difference of the Percent Spliced In (dPSI) index values, indicating the strength and direction of AS events that differ between transient lncRNA expression and the GFP control. The NSRa bound lncRNAs *1-NSRa* (AT2G15292) and *2-NSRa* (AT2G18735) had the strongest impact on AS, being associated with over 350 differential AS (DAS) events (Figure 2A). We therefore decided to focus our functional analysis on these two lncRNAs. *1-NSRa* was an uncharacterized lncRNA and we named it *ACHLYS*. The candidate *2-NSRa*, identified in our transient expression screen, has also been described as *FLAIL* and shown to influence gene expression and AS of flowering-related genes (Jin et al., 2023). The GRP7 bound lncRNAs with the biggest effect on AS were *1-GRP7* and *2-GRP7* with more than 250 DAS events. The most frequent splicing events detected were intron retention (30-40%) and alternative 3’- (24-34%) and 5’-SS (12-20%) (Figure S1D).

Comparison between the DAS genes and the differentially expressed genes (DEGs) showed that only a small fraction of DAS genes overlap with DEGs (*FLAIL* 22%, *1-GRP7* 11% and *2-GRP7* 27%) (Figure S1C).

Notably, *ACHLYS* has a greater effect on AS (378 DAS genes) than on steady-state mRNA levels (293 DEGs) (Figure 2A). Whereby only 1% of the DAS genes are differentially expressed (Figure S1C).

The AS profiles of the eleven lncRNAs, (three interacting with NSRa, six with GRP7 and two with none of the two SFs), allowed us to test if the lncRNAs affected unique or overlapping DAS genes. Therefore, we intersected the DAS genes of all eleven lncRNAs against each other. The upset graph shows the first 7 intersections with the highest intersection size of the lncRNA induced DAS genes (Figure 2B). 34% of the DAS genes affected by *FLAIL* were unique, 26% for *ACHLYS*, 23% for *1-GRP7* and 19% for *2-GRP7*, indicating that the lncRNAs, despite interacting with the same SF, affect mainly different individual DAS genes. No major overlaps between DAS genes regulated by the six GRP7 bound lncRNAs were detected. In contrast, the two NSRa bound lncRNAs *ACHLYS* and *FLAIL* share 19% and 21% of DAS genes, respectively. Surprisingly, *ACHLYS* and *FLAIL* do not only share DAS in the same genes but also 86 identical splicing events, making up around 15% of all *ACHLYS* and *FLAIL* induced AS events. Moreover, calculated dPSI values of *ACHLYS* and *FLAIL* induced DAS events are strongly positively correlated (R^2^ = 0.94), suggesting that they affect these DAS events in the same direction (Figure 2C).

To validate individual splicing events, we perform qPCR splicing analysis. We first quantified for each DAS gene the RNA level of a shared exon, as an internal normalization. Second, we used the exon–exon junction spanning primers to quantify the spliced mRNA and third, using primers detecting only the AS event (unspliced). We confirmed for three selected AS events the reduced intron retention after transient *ACHLYS* or *FLAIL* expression (Figure 1D).

In summary, we find that lncRNAs interacting with the same SF mainly affect AS of unique mRNAs. Nevertheless, AS of a subgroup of mRNAs is affected both by *ACHLYS* and *FLAIL*, in the same way, indicating that they may regulate this AS event through a shared mechanism.

### Characterization of the *ACHLYS* locus and its expression

*ACHLYS* is a 1253 nt long mono exonic gene annotated as lncRNA in TAIR11 (www.arabidopsis.org). Its coding potential is predicted to be low with a score of -0.41 by CNIT (-1 non-coding to 1 coding) and 0.04 by CPC2 (0 non-coding to 1 coding) (Kang et al. 2017; J.-C. Guo et al. 2019) (Figure S1A). Based on Ribosome profiling of total seedlings (H.-Y. L. Wu et al. 2024), *ACHLYS* is not associated with ribosomes, and its annotated transcriptional start site (TSS) is in agreement with TSS-seq results (Nielsen et al. 2019) (Figure 3A). *ACHLYS* contains a 99 bp SINE fragment (AT2TE27080) at its 5’-end and two RC/Helitrons are located upstream (AT2TE27085, 1483 bp and AT2TE27090, 46 bp) of the TSS (Figure 3A).

**Figure 3:**
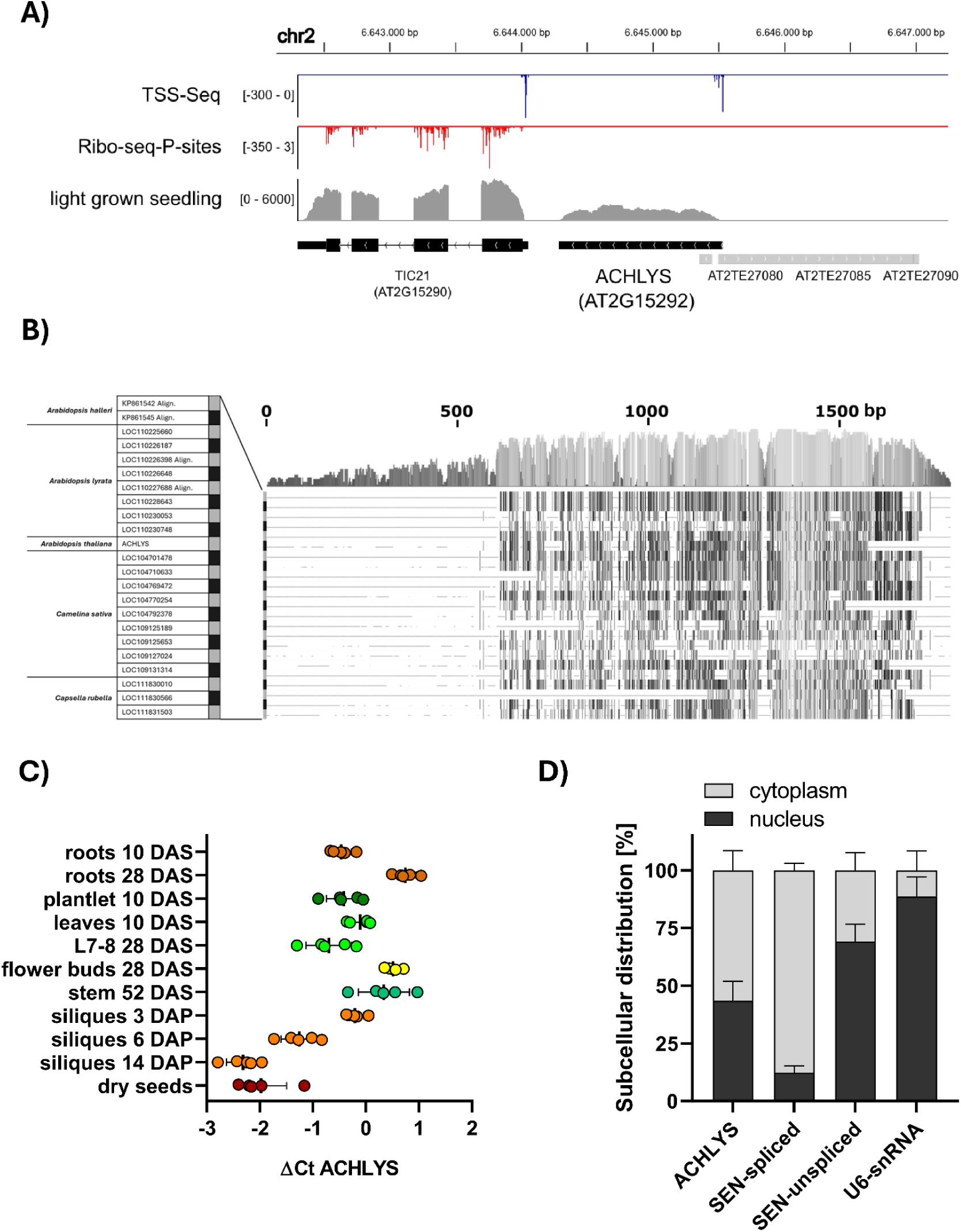
Characterization of the *ACHLYS* lncRNA. **A)** *ACHLYS* locus and neighboring genes. Lines indicate introns; small boxes, UTRs; large boxes, CDS; and grey boxes, transposable elements. Transcriptional start sites from (TSS)-seq are displayed in blue (Nielsen et al. 2019), Ribosome Profiling in red and coverage plot of light grown seedling in grey (H.-Y. L. Wu et al. 2024). **B)** Sequence alignment of *ACHLYS* (*A. thaliana*) and *ACHLYS*-like sequences in other Brassicaceae species. Grey scale and bar graph indicate nucleotide conservation along all aligned sequences. **C)** *ACHLYS* RNA levels normalized to PP2A in different organs and time points. DAS (days after sowing), DAP (days after pollination), L7-8: seventh and eighth leaf. **D)** Subcellular localization of *ACHLYS* estimated by qPCR after fractionating cytoplasm and nucleus. The SEN-spliced mRNA is considered a cytoplasmic control and SEN-unspliced and U6 snRNA as positive control for nuclear localization. The subcellular distribution was calculated as the percentage of 2^ΔCt-cytoplasm^+ 2 ^ΔCt-nucleus^. *ACHLYS* is enriched in the nuclear fraction compared to the SEN mature mRNA although not as the SEN mRNA precursor or the UsnRNA.

We examined whether *ACHLYS* is conserved in other plant species using *ACHLYS* as a query for the NCBI Nucleotide BLAST against refseq_rna of *Brassicaceae*. 23 sequences were highly similar (68-86 Percent Identity with coverage > 30%,) to *A. thaliana ACHLYS*, particularly at its 3’-half. While in *A. thaliana* only one copy of *ACHLYS* was found, other species encode multiple copies. In *A. lyrata* eight sequences were similar to *ACHLYS*, in *A. halleri* two, *Camelina sativa* nine and *Capsella rubella* three (Figure 3B). *C. sativa* and *A. thaliana* shared the last common ancestor 9 million years ago according to TimeTree 5 (Kumar et al. 2022).

Next, we compared the expression profile of *ACHLYS* during the plant development by qPCR using different tissues. Globally, *ACHLYS* is expressed throughout development at RNA levels comparable to the housekeeping gene *PP2A*, with the highest expression observed in 28-day-old roots and lowest in siliques 14 days after pollination, showing an 8-fold difference between these two points (Figure 3C). In a complementary approach, we exploited more than twenty-eight thousand publicly available *A. thaliana* RNAseq datasets (H. Zhang et al. 2020). These data confirmed that *ACHLYS* is stably expressed in different tissues, and no specific developmental stage or condition showed a major variation in expression (Figure S2A). The median *ACHLYS* FPKM (fragments per kilobase of transcript per million mapped reads) shows 8-times higher expression than *ASCO*, 2.7-times higher than *FLAIL*, similar expression as NSRa, and 4-times less than GRP7. Considering that *ACHLYS* is regulated during LRF (Figure S1A), we used publicly available single-cell RNAseq data to visualize its expression levels across different root cell types. *ACHLYS* expression is highest in xylem cells but detectable in many cell types. The largest differences between related cell types were observed between trichoblast (0.25) and atrichoblast cells (1.75), as well as between young epidermis (2.0) and mature epidermis (0.75) (Figure S2B). Notably, this data confirmed similar expression levels of NSRa and *ACHLYS* and proposes that a change in *ACHLYS* expression can be at the basis of a potential biological function in root development.

LncRNAs that regulate AS are expected to be enriched in the nucleus (Bejerano et al. 2004; Hutchinson et al. 2007; Gonzalez et al. 2015). Therefore, we fractionated the cytoplasm and nucleus of total seedlings, extracted the RNA and performed qPCR to estimate *ACHLYS* subcellular localization. Results indicate that 43% of *ACHLYS* transcripts are located in the nucleus. While this is lower than the nuclear enrichment observed for the pre-mRNA control (69% SEN1-unspliced) and spliceosomal small nuclear RNA U6 (snRNAU6) (89% U6snRNA), it is substantially higher than that of a typical mature mRNA (12% SEN1-spliced), suggesting that *ACHLYS* may undergo nuclear retention or have a nuclear-related function (Figure 3D). Taken together, our data show that *ACHLYS* is a highly abundant lncRNA, enriched in the nucleus and expressed uniformly across Arabidopsis tissues.

### Deregulation of the lncRNA *ACHLYS* affects alternative splicing

To study if *ACHLYS* deregulation affects AS in stable plants, we first isolated two T-DNA insertion lines (Figure S3A). The first T-DNA insertion is located 658 bp downstream from the *ACHLYS* TSS in the gene locus (*ACHLYS-1*). qPCR analysis detected just 2 times lower expression levels of a truncated transcript (Figure S3B). The second T-DNA (*ACHLYS-2*) is inserted in the promoter (-228 bp from TSS) resulting in 2 times higher *ACHLYS* levels (Figure S3B). Since both lines showed minor changes in *ACHLYS* expression, only a single T-DNA insertion mutant at a 3’ position was available and the T-DNA insertion deregulated neighboring genes of *ACHLYS* (Figure S3C), we did not pursue any further phenotypical characterization of these lines.

To deregulate *ACHLYS* levels, we generated two independent RNAi knockdown (KD-*ACHLYS*) and 35S Cauliflower Mosaic Virus promoter overexpression (OE-*ACHLYS*) lines. KD-*ACHLYS*-1 and -2 showed an 8-fold decrease, while OE-*ACHLYS*-1 and -2 lines exhibited a 32-fold increase in *ACHLYS* expression compared to the wild type (Figure 4A). To assess the effect of low and high *ACHLYS* levels on AS and transcript levels, we performed stranded poly(A)-RNAseq on pools of 10-day-old KD-*ACHLYS*-1 and OE-*ACHLYS*-1 seedlings. In KD-*ACHLYS* plants, 61 genes were differentially expressed (Figure 4B), and gene ontology analysis revealed a more than 10-fold enrichment for genes involved in hypoxia and ethylene signaling (Table S1). In OE-*ACHLYS* plants, only seven DEGs were found, including *ACHLYS* itself (Figure S4B). No DEGs overlapped between OE-*ACHLYS* and KD-*ACHLYS* lines (Figure 4B). Thus, compared to other well-characterized lncRNAs (Ariel et al. 2014; Rigo et al. 2020; Jin et al. 2023), *ACHLYS* deregulation has a limited impact on steady state poly(A) RNA levels of specific transcripts.

**Figure 4:**
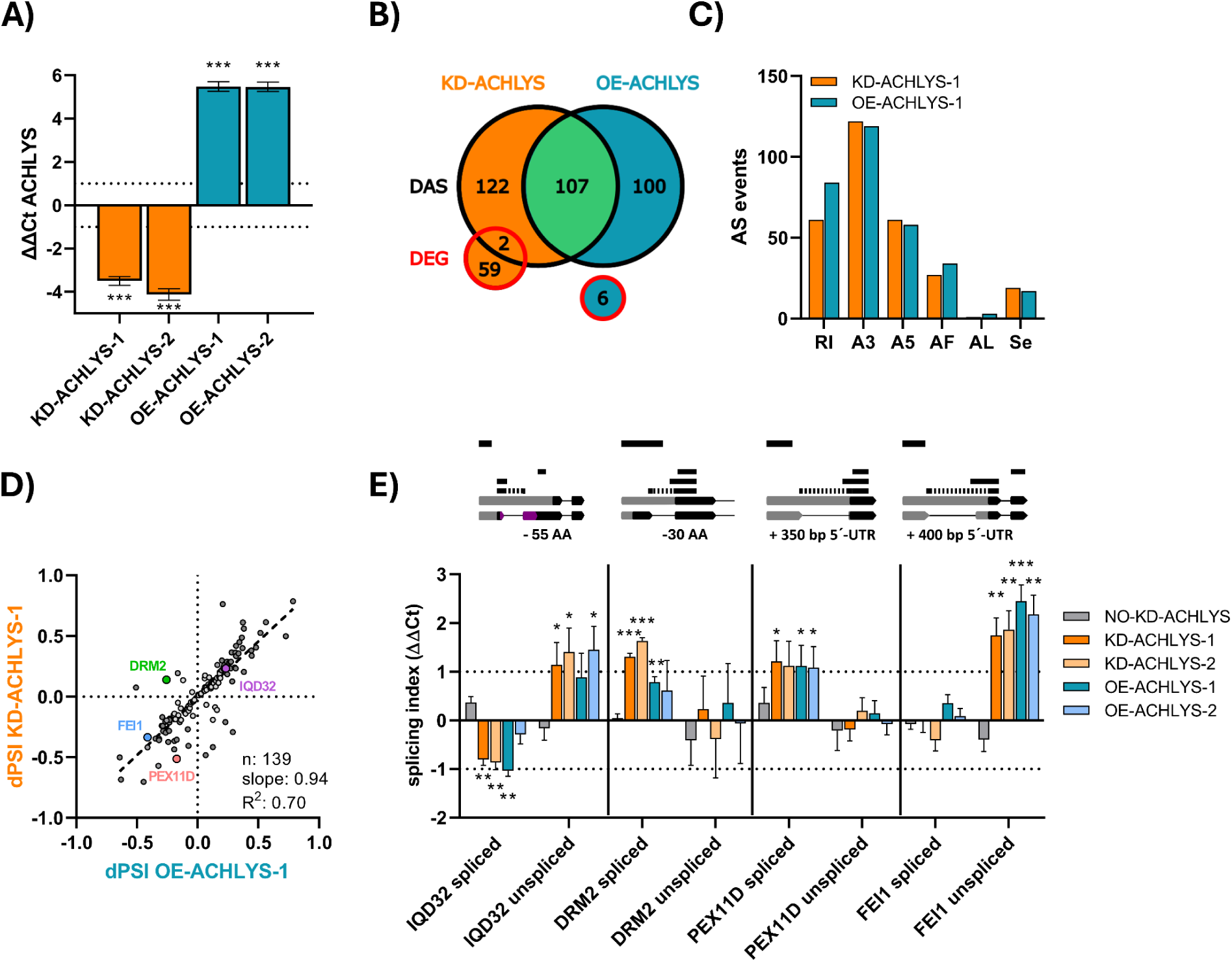
*ACHLYS* deregulation affects alternative splicing in plants. **A)** *ACHLYS* RNA levels in 10-day-old 35S:RNAi-*ACHLYS* (KD-*ACHLYS*) and 35S:*ACHLYS* (OE-*ACHLYS*) seedlings. ΔΔCt is the mean difference (ΔCt mutant minus ΔCt control, using PP2A as housekeeping gene, comparing 3 to 4 biological replicates) and standard error of mean difference is shown. Dotted lines indicate two-fold change (unpaired, two-tailed T-test, ***p ≤ 0.001). **B)** Venn diagram of differential alternatively spliced (DAS, black bordered) genes determined by SUPPA2 and differentially expressed genes (DEG, red bordered) based on DEseq2 in 10-day-old KD- and OE-*ACHLYS* seedlings. **C)** Number of different alternative splicing events detected in KD- and OE-*ACHLYS*. Splicing events are classified as retained introns (RI), alternative 3’ splice (A3), alternative 5’ splice (A5), alternative first exon (AF), and skipped exon (SE). **D)** Positive correlation between delta percent spliced in (dPSI) of shared AS events detected in KD-and OE-*ACHLYS*. Alternative splicing events validated by qPCR are highlighted. Grey dots mark dPSI between -0.2 and 0.2. Number of DAS events (n), slope and R2 of the linear regression are shown. **E)** Confirmation of individual AS events in 10-day-old seedlings of two independent KD-and OE-*ACHLYS* lines and as an additional control, we use an *ACHLYS* RNAi line carrying the transgene but with no effect on *ACHLYS* RNA levels (No knock down or NO-KD-*ACHLYS*). ΔΔCt as mean difference (ΔCt was calculated as Ct-spliced/unspliced minus Ct-constitutive exon and ΔΔCt as ΔCt mutant minus ΔCt wild type comparing 3 to 4 biological replicates) and standard error of mean difference. Dotted lines indicate two-fold change (unpaired, two-tailed T-test, *p ≤ 0.05, **p ≤ 0.01, ***p ≤ 0.001). Upper schemes are a zoom on gene locus of detected alternative splicing events. Exons are shown as bars (URTs grey and CDS black) and introns as lines. qPCR products are shown in black lines and intron spanning primers as dotted lines. Upper black lines represent a scale of 100 bp.

Nevertheless, analysis of splicing patterns using SUPPA2 revealed that KD-*ACHLYS*-1 exhibits 231 DAS genes, compared to 61 DEG genes (Figure 4B, Figure S4A,C). Similarly, we detected 207 DAS genes in contrast to the 7 DEG for OE-*ACHLYS*-1 (Figure 4B, Figure S4B,D). This data resemble the tendency observed after the transient overexpression of *ACHLYS* in leaves, where we also detected more DAS than DEG genes (Figure 2A,B). The main AS events for KD-and OE-*ACHLYS* are alternative 3’-SS (122 and 119) followed by intron retention (61 and 84) and alternative 5’-SSs (61 and 58) (Figure 4C).

Notably, 139 identical splicing events were detected in OE-*ACHLYS* and KD-*ACHLYS*, which represents more than 40% of all detected splicing events. Surprisingly, the dPSI of the shared DAS events between KD- and OE-*ACHLYS* lines are mainly positively correlated (100 out of 114, Figure 4D). We performed gene ontology analysis on these common DAS events showing a dPSI larger than 0.2 or smaller than -0.2 at least in one of the lines. The biological function GO-terms “alternative mRNA splicing via spliceosome“ (FDR = 5.2*10^-3^), “meristem growth“ (FDR = 3.5*10^-2^) and “cellular component organization“ (FDR = 3.5*10^-2^) were enriched, as well as the cellular compartment GO terms “peroxisome“ (FDR = 2.8*10^-3^), “cell-cell junction“ (FDR = 2.9*^-6^) and “plasma membrane“ (FDR = 9.4*10^-3^) (Table S2).

To confirm individual splicing events, we selected intron retention events in four genes that were deregulated in both KD- and OE-*ACHLYS* lines. We tested these splicing events in two independent KD- and OE-*ACHLYS* lines by qPCR and compared them to wild type plants. As an additional control we used a RNAi-*ACHLYS* line, in which *ACHLYS* levels are similar to wild type (NO-KD-*ACHLYS*) (Figure S3D).

In KD- and OE-*ACHLYS* the retention of the first intron of IQD32 (IQ-domain 32 protein, AT1G19870) leads to a longer 5’-UTR and a premature stop codon. Nevertheless, an annotated alternative translation start site could potentially lead to the translation of a protein lacking the 44 amino acid N-terminal IQ domain. The IQD32 can bind microtubules and is involved in calcium signaling (Bürstenbinder et al. 2017). In KD-*ACHLYS* lines, the first intron of DRM2 (dormancy associated gene 2, At2g33830) is spliced more efficiently, resulting in the accumulation of an mRNA isoform that encodes an additional 30 amino acids at the N-terminus. These amino acids are potentially absent in the mRNA isoform with the retained intron due to an alternative translation start site.

DRM2 is a repressor of pathogen response by suppressing ROS accumulation and local callose deposition (Roy et al. 2020). *ACHLYS* deregulation leads to higher levels of a PEX11D (PEROXIN11D, AT2G45740) mRNA isoform with shorter 5’-UTR due to higher levels of spliced mRNA.PEX11D is located in peroxisomes and controls peroxisome proliferation (Orth et al. 2007). *ACHLYS* deregulation leads to an increased proportion of FEI1-mRNAs (AT1G31420) containing a 5’-UTR extension of around 400 nt due to an intron retention event (Figure 4E). FEI1 is a leucine-rich repeat receptor kinase regulating cell wall elongation in the context of root development (Xu et al. 2008).

Considering that *ACHLYS* and *FLAIL* both interact with NSRa, we explored whether KD-*ACHLYS* share splicing events with the *FLAIL* T-DNA insertion line *flail3*, which was characterized as a *FLAIL* knockout (Jin et al. 2023). Analysis of public *flail3* RNAseq data and its intersection with KD-*ACHLYS* data showed that they share 21% (66) of splicing events. Strikingly, 51 of the shared events are negatively correlated and only 13 show a positive correlation (Figure S4E). This data indicate that *ACHLYS* knockdown and *FLAIL* knockout have a predominantly opposite effect on shared splicing events.

### *ACHLYS*- and *FLAIL*-dependent alternative splicing events are locally enriched in NSRa binding sites

To understand whether the splicing events detected after transient *ACHLYS* expression or in KD- and OE-*ACHLYS* lines are potentially regulated by NSRa, we set out to map *in vivo* NSRa binding sites transcriptome wide by iCLIP. We used a *nsra* mutant line complemented with pNSRa:NSRa-GFP (NSRa-GFP; Bazin et al. 2018) and followed the iCLIP protocol described by Lewinski et al. 2024. To increase NSRa-GFP expression and accordingly bound RNA as input for NSRa-iCLIP, we exposed 14-days-old NSRa-GFP seedlings to 38°C for 3 h before crosslinking.

The NSRa-iCLIP identified 20,649 reproducible RNA binding sites with a width of 3 nt detected in at least 2 out of 3 biological replicates (Figure S5A). Of these, 36% mapped to protein-coding transcripts, 7% to annotated non-coding transcripts. Within protein-coding genes, binding sites were more prevalent in 3’-UTRs, 5’-UTRs and introns relative to the length of the feature in the genome (Figure S5B). A more in-depth analysis of peak-score distributions for the same protein-coding targets revealed that higher scores were associated predominantly with 3’-UTRs and introns (Figure S5C). This trend was promoted for 3’-UTRs by normalizing to the relative size of the genomic features with the addition that 5’-UTRs follow this trend towards higher peak scores (Figure S5D). Therefore, the highest scored NSRa-GFP binding sites resided mostly in the UTRs, higher than expected in intronic regions and are depleted from CDS.

To identify the NSRa binding motif we used two *de novo* motif discovery approaches. First, using STREME two motifs with C-duplet were found to be in direct vicinity of the NSRa-GFP binding sites with high coverage of 60% and 37% (Figure 5A). Second, we counted occurrences of trimers in the binding sites of 3 nt width, which span the estimated binding site width of NSRa-GFP. The top counted trimers were uCc, uUu and cCc where the capital letter denotes the center nucleotide of the binding site (Figure S5E). The application of z-scoring using a uniform background model in the identical region of the mapped binding site with 100 repetitions revealed that the most enriched trimers besides uUu are cytosine duplets or triplets. These findings are in line with the independent motif discovery by STREME. Therefore, the cytosine duplets and triplets are core drivers for the NSRa-GFP binding site choice.

**Figure 5:**
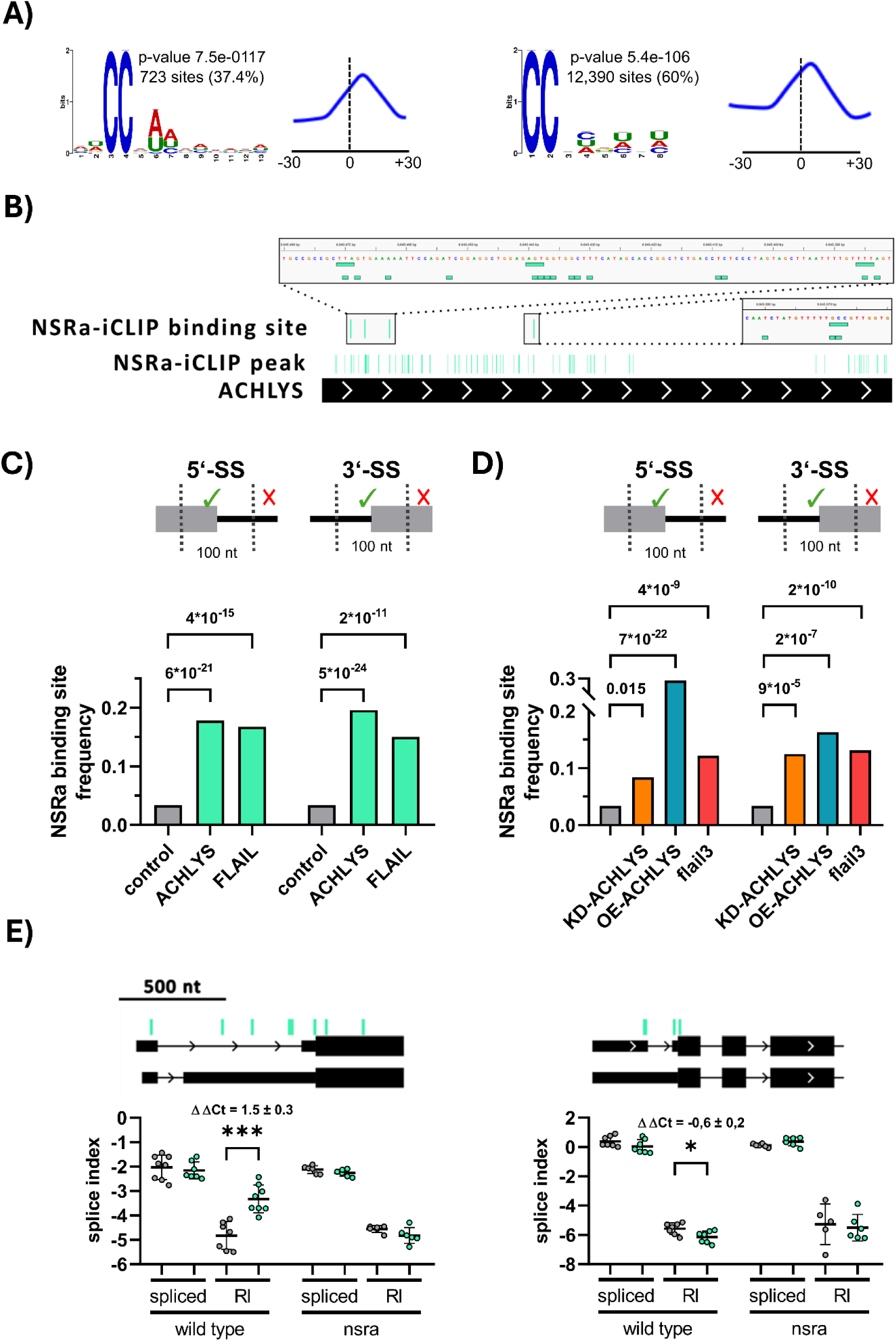
NSRa-iCLIP revealed that *ACHLYS* and *FLAIL* dependent DAS events are enriched in NSRa binding sites. **A)** Significantly enriched motifs determined by STREME. The blue line illustrates the density of the motif in an area of 30 nucleotides up- and downstream of the binding site peak of NSRa-iCLIP. **B)** NSRa-iCLIP peaks based on peak calling of crosslinking site and statistically significant binding site based on enriched peaks at *ACHLYS* transcript. **C)** Percentage of intron retention events with NSRa-iCLIP binding sites in a 100 nt window centralized at the 5’- and 3’-splice sites (SS). Intron retention events were detected after transient expression of *ACHLYS* and *FLAIL* or **D)** in *ACHLYS* knockdown (KD), overexpression (OE) and *flail3* knockout lines (right), compared to frequency of intron retention event with NSRa-iCLIP binding sites of all annotated intron retention events (grey). Statistically significance was tested by Hypergeometric test and phyper values are displayed. **E)** RNA levels of spliced and intron-retained (RI) isoform after transient expression of *ACHLYS* (green) compared to GFP (grey) in wild type and *nsra* knockout mutant. Dots represent individual biological replicates (n = 6-7, unpaired, two-tailed T-test, *p ≤ 0.05, **p ≤ 0.01, ***p ≤ 0.001).

After genome wide description of NSRa binding sites we focused our analysis on lncRNA dependent AS events. We identified three NSRa binding sites in the 5’-region and one in the center of *ACHLYS* (Figure 5B), four NSRa binding sites in the 3’-half of *FLAIL* and two at the antisense *FLAIL* transcript identified by Jin et al. 2023. This suggested that the interaction between NSRa and the lncRNAs previously determined by RIP-seq were due to direct binding (Bazin et al. 2018).

The iCLIP data allowed us to address if lncRNA-dependent AS events are linked to NSRa binding. First, we tested whether NSRa-iCLIP peaks are enriched in the *ACHLYS* and *FLAIL* induced DAS genes after transient expression in *A. thaliana* leaves. If NSRa binding sites were randomly distributed, we would expect approximately 8.3% NSRa-bound RNAs across the transcriptome. Interestingly, 32–36% of *ACHLYS*- and *FLAIL*-dependent DAS genes exhibited NSRa-iCLIP binding sites (Figure S6A).

The NSRa-iCLIP data also allowed us to assess the positional relationship between NSRa binding sites and different types of AS events. We therefore analyzed intron retention, alternative 5’- and 3’-SS events and quantified the fraction of lncRNA-dependent AS events with NSRa binding sites within a 100 nt window centered on the 5’- or 3’-SS (Figure S6C-D).

Notably, a significant enrichment was observed for intron retention events: approximately 15% of *ACHLYS*- and *FLAIL*-dependent intron retention events contained at least one NSRa binding site near the 5′- or 3′-splice site, compared with 3.4% of all annotated intron retention events (Figure 5C). No comparable enrichment was detected for other AS types. These results indicate that NSRa binding is specifically associated with *ACHLYS*- and *FLAIL*-dependent intron retention events. Next, we performed the same analysis for the DAS events identified in KD-and OE-*ACHLYS* plant lines and *flail3* (Jin et al. 2023). Similar to the results obtained for the transiently expressed lncRNAs, 23-31% of the DAS RNAs regulated by *ACHLYS* or *FLAIL* are bound by NSRa (Figure S6B). Intron retention events with NSRa binding sites around the 5’- or 3’-SS are with 8-29% statistically significantly enriched in the stable KD-, OE-*ACHLYS* and flail3 lines (Figure 5D). This analysis of NSRa binding sites showed that *ACHLYS*- and *FLAIL*-dependent AS events, in the transient expression experiment and in stable plant lines, are enriched in NSRa binding sites, both at gene level and, more specifically, in direct proximity to the affected AS events. This result suggests that *ACHLYS* and *FLAIL* regulate AS through modulation of NSRa activity, particularly on intron retention events.

To test this hypothesis, we selected AS events identified after transient *ACHLYS* expression that also exhibit multiple NSRa binding sites near the splicing event. We focused on an intron retention event at 5’-UTR for glutamate dehydrogenase 2 (GDH2), which displays 8 significant NSRa binding sites, and on another one for translation initiation factor SUI1 5’-UTR, which exhibits 4 significant NSRa binding sites.

After transient expression of *ACHLYS* in the seventh leaf of wild type plants, we observed more intron retention for GDH2 (ΔΔCt = 1.5 ± 0.3) and less for SUI1 (ΔΔCt = -0.6 ± 0.2) compared to controls (Figure 5E). In the same experiment, we transiently overexpressed *ACHLYS* in *nsra* knockout plants and showed that the levels of the individual GDH2 and SUI1 isoforms are similar to the GFP control, strongly supporting that individual *ACHLYS*-induced splicing events required NSRa function.

### *ACHLYS* deregulation and *FLAIL* knockout lines exhibit an opposite root phenotype

After establishing *ACHLYS* as an AS regulator, we wanted to understand its biological function. *ACHLYS* and NSRa expression is differentially regulated during LRF (Figure S1A). Also, *nsra/b* double mutants show a reduced number and length of auxin-induced lateral roots (Bardou et al. 2014). Therefore, we performed root phenotyping of KD- and OE-*ACHLYS* plants. We measured the main root length and lateral root length of 14-day-old seedlings in three independent experiments. In two independent KD-*ACHLYS* and OE-*ACHLYS* lines we quantified a slight but statistically significant shorter main root length of -5% and -7%, respectively (Figure S7A). The lateral root density is slightly reduced in deregulated *ACHLYS* lines and only significant for KD-*ACHLYS*-2 and OE-*ACHLYS*-1 (Figure S7B). The OE-*ACHLYS* lines also show significantly shorter lateral root lengths (-24%) (Figure S7C) and consequently shorter total root system length (-16%) compared to wild type (Figure S7D). KD-*ACHLYS* plants have slightly shorter lateral root lengths (Figure S7C) and total root system lengths, only statistically significant for KD-*ACHLYS*-2 (Figure S7D).

Considering that *FLAIL* is also deregulated during LRF (Figure S1A) and interacts with NSRa, we performed root phenotyping on two *FLAIL* CRISPR deletion lines (*flail1* and *flail2*) and a T-DNA line (*flail3*), which perturb the *FLAIL* locus in *cis* (Jin et al. 2023). Three independent experiments showed that all three *FLAIL* knockout lines exhibited longer main root length (Figure S7E), and no effect on lateral root density compared to wild type (Figure S7F). Moreover, we quantified the main root length of the *flail3* line complemented with p*FLAIL*:*FLAIL*. Quantification of *FLAIL* RNA levels showed that *flail3* p*FLAIL*:*FLAIL*-1 complements *FLAIL* RNA to wild type levels, while *flail3* p*FLAIL*:*FLAIL*-2 show reduced *FLAIL* levels similar to *flail3* mutant (Figure S7G). As control, we used a *flail3* line expressing the GUS transgene under the control of the 35S promoter (*flail3* OE-GUS). In three independent experiments the *flail3* OE-GUS and *flail3* p*FLAIL*:*FLAIL*-2 showed a longer main root length compared to wild type and the fully complemented line *flail3* p*FLAIL*:*FLAIL*-1 (Figure S7H). These experiments suggest that perturbation of the *FLAIL* locus through T-DNA or CRISPR deletion increases main root lengths.

To link the observed root phenotypes and AS regulation, we intersected our lncRNA dependent DAS genes with GO term lists related to “root” and found that 6% KD-, OE-*ACHLYS* and *flail3* DAS genes could be linked to root development. The effect of individual AS events on the function of these genes still needs to be elucidated. Furthermore, we intersected the DAS event list of the LRF kinetics with lncRNA dependent AS events and identified that up to one third of the same AS events could be detected in KD- (34%), OE-*ACHLYS* (28%) and *flail3* (29%). These DAS genes potentially related to root development include genes for which we characterized AS events by qPCR (Figure 4E). For instance, the double mutant of FEI1 (fei1/2) exhibits shorter main root length (Xu et al. 2008). Also, the knockout of IQD32 lacks root elongation as response to sulfur scavenging (Grubb et al. 2025) and its AS is regulated during our LRF kinetics.

Taking together, *ACHLYS* deregulation impairs root development and exhibits an opposite effect on main root length compared to plants with a *FLAIL* knockout. This opposite phenotype was also previously observed molecularly for common AS events potentially linked to root development of *ACHLYS* and *FLAIL* knock down and knockout, respectively.

### High *ACHLYS* levels promote *in vitro* and *in vivo* biomolecular condensate formation of NSRa

NSRa subcellular localization is in the nucleus and mainly in Nuclear speckles (Bardou et al. 2014). It was shown in transient expression assay that the *Medicago truncatula* lncRNA ENOD40 can decoy the NSRa homologue MtRBP1 from Nuclear speckles to cytoplasmic granules depending on its RNA recognition motif (RRM) (Campalans et al. 2004). Moreover, other lncRNAs are reported to change the association of SFS with nuclear biomolecular condensates *in vivo* (Tripathi et al. 2010; Z. Zhang et al. 2016). Many SFs undergo phase separation *in vitro*, which could be quantified by turbidity measurements (Huang et al. 2021). Notably, RNA molecules are known to promote *in vitro* phase separation of RNA binding proteins, including SFs (De Vries et al. 2024). As *ACHLYS* interacts with NSRa, we decided to assess the ability of the lncRNA *ACHLYS* to promote NSRa condensate formation *in vitro*. Hence, we first generated an RNA-binding–deficient mutant version of NSRa-(FD) by substituting a single highly conserved Phenylalanin (F184) within the RRM domain to Aspartic acid (D), specifically in the β-sheet containing the ribonucleoprotein domain 1 (RNP1) previously described as the principal determinant of RNA recognition (Cléry et al. 2008; Daubner et al. 2013) (Figure 6A). We then validated the loss of RNA-binding activity using RNA Electrophoretic Mobility Shift Assay (RNA-EMSA) assays. For this, we purified the GST-tagged RRM domain of NSRa from bacteria both the wild type (NSRa-RRM) or the RNA binding mutant (NSRa-RRM-(FD)) and incubated it with a 22-nt RNA probe corresponding to the *ACHLYS* region showing the highest NSRa iCLIP peak (First 22 nt in Figure 5B covering the first NSRa-iCLIP binding site), alongside a scrambled version of that region as a negative control probe. The absence of probe shift with the mutant version allows us to conclude that the FD mutation abolished RNA binding of NSRa-RRM *in vitro* (Figure 6B). We next examined the phase-separation behavior of these RRM domains. Recombinant NSRa-RRM (WT) was mixed with increasing concentrations of full-length *in vitro*–transcribed *ACHLYS* RNA, and condensate formation was monitored by turbidity measurements at OD₄₀₀, as previously described (Chen et al. 2023). Intermediate concentrations of *ACHLYS* markedly stimulated NSRa-RRM condensate formation, whereas no such effect was observed with the NSRa-RRM-(FD) mutant, indicating that *ACHLYS*-driven phase separation strictly depends on NSRa’s RNA-binding capacity (Figure 6C). In addition, we assessed whether this condensate-promoting activity was specific to *ACHLYS* by testing other *in vitro* transcribed RNAs of comparable length and composition. We included the antisense version of *ACHLYS* (*ACHLYS*as) and a 1.3-kb fragment of the GUS coding sequence, which matches the size of *ACHLYS*. Under the same assay conditions, all three RNAs promoted NSRa-RRM condensate formation *in vitro* to a similar extent (Supplementary Fig. S8A).

**Figure 6:**
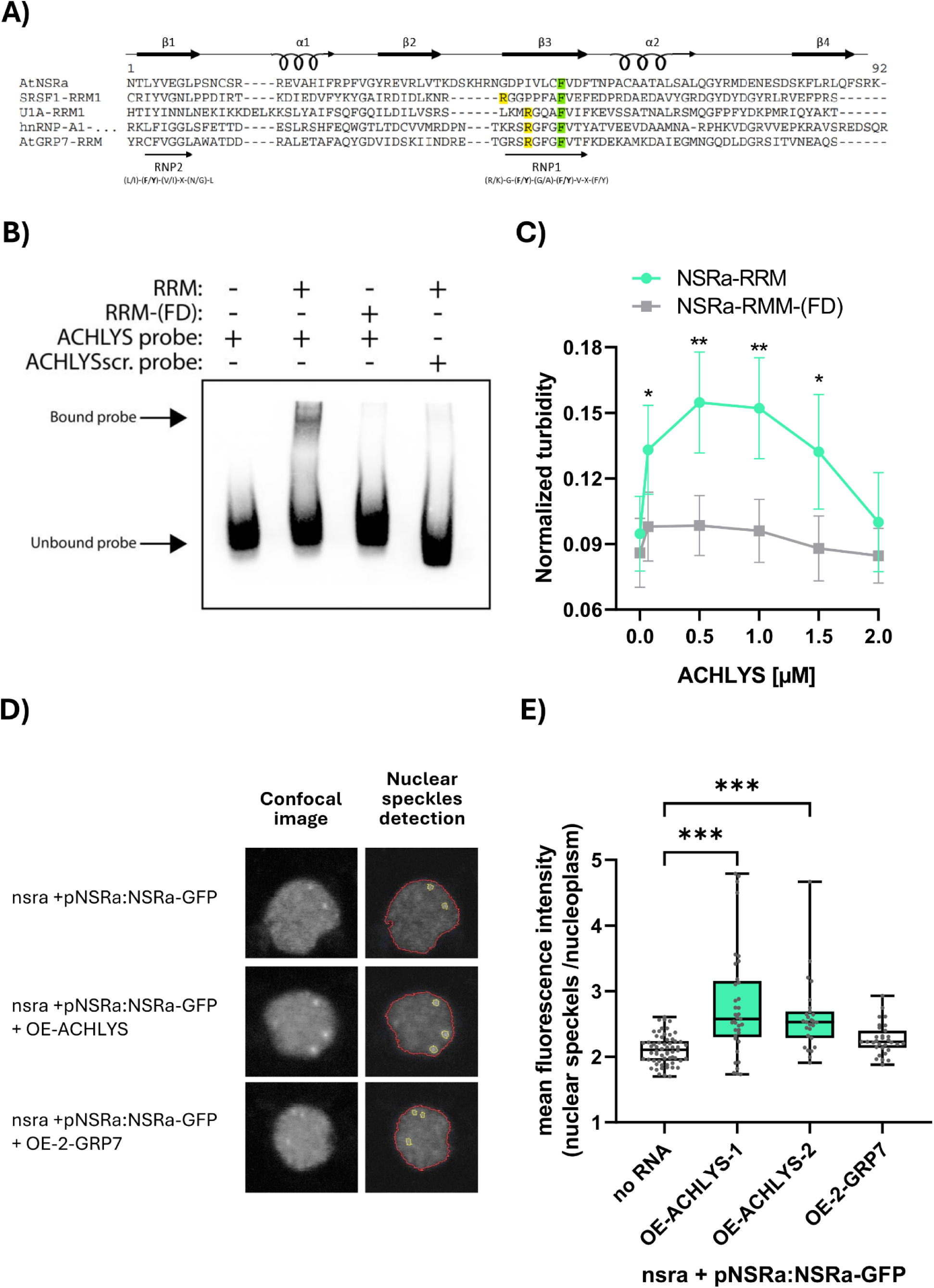
High *ACHLYS* levels increase NSRa concentrations in Nuclear speckles. **A)** Amino acid alignment of four splicing factor RRMs including NSRa. Essential regions for RNA binding the Ribonucleoprotein (RNP) 1 and 2 are marked with arrows. To generate a potential NSRa RNA binding mutant, the central Phenylalanin (F184) within the RNP1 labeled in green was replaced by an Aspartic acid (D). **B)** Electrophoretic Mobility Shift Assay (EMSA) of NSRa-RRM or NSRa-RRM-(FD) with biotinylated 22-nt RNA probe corresponding to the *ACHLYS* sequence described in M&M or scrambled probe with the same nucleotide composition but different order (*ACHLYS*scr.). **C)** *In vitro* phase separation of NSRa-RRM or NSRa-RRM-(FD) (6µM) in the presence of increasing concentrations of *in vitro* transcribed *ACHLYS* (1234 nt). Mean and 95% confidence interval (n = 13-20, multiple unpaired, two-tailed T-test, *p ≤ 0.05, **p ≤ 0.01, ***p ≤ 0.001). **D)** Representative pictures showing automatic detection of nucleus (red outline) and Nuclear speckles (yellow outline) using CellProfiler in epidermis cells of 6-day-old root meristem in *nsra* pNSRa:NSRa-GFP plants stably overexpressing lncRNAs under the 35S promoter. **E)** Mean NSRa-GFP fluorescence intensity of all Nuclear speckles detected from one cell normalized to NSRa-GFP fluorescence intensity in the nucleoplasm in *nsra* pNSRa:NSRa-GFP plants stably overexpressing lncRNAs under the 35S promoter (n = 27-62). Box-and-whisker plots (median, first and third percentiles, whiskers min-max percentile, dots represent biological replicates), one-way analysis of variance (ANOVA) and Tukeýs honestly significant difference (HSD) post-hoc test, ***p ≤ 0.001.

After establishing that *ACHLYS* potentially affects AS through its interaction with NSRa and that direct interaction of NSRa with *ACHLYS* promotes *in vitro* phase separation, we wondered if *ACHLYS* could modulate *in vivo* NSRa association with Nuclear speckles. To mimic the *in vitro* phase separation experiment *in planta,* we stably transformed nsra mutant lines complemented with pNSRa:NSRa-GFP (WT-NSRa-GFP) with constructs overexpressing *ACHLYS* (OE-*ACHLYS*-NSRa-GFP) and the GRP7 bound lncRNA *2-GRP7* (OE-*2-GRP7*-NSRa-GFP) and observed NSRa-GFP localization by confocal microscopy in root tips of 6-day-old seedlings.

In OE-*ACHLYS*-NSRa-GFP lines, we observed NSRa-GFP localization in the nucleus and in Nuclear speckles and no global effect on NSRa-GFP subcellular localization compared to WT-NSRa-GFP. However, the NSRa signal appeared more intense within Nuclear speckles. We used the image analysis software CellProfiler (Carpenter et al. 2006) to quantify the NSRa signal within Nuclear speckles. The program automatically detects and quantifies the signal intensity of NSRa-GFP in Nuclear speckles and the nucleoplasm. We then compared the Nuclear speckle signal intensity in WT-NSRa-GFP, two OE-*ACHLYS*-NSRa-GFP lines and one OE-*2-GRP7*-NSRa-GFP by normalizing the mean intensity of Nuclear speckles to the mean intensity of the nucleoplasm. Strikingly, only *ACHLYS* overexpression leads to a significant increase of NSRa-GFP signal within Nuclear speckles (Figure 6D), suggesting that the interaction of *ACHLYS* with NSRa could increase its recruitment to Nuclear speckles.

To determine whether the observed differences in accumulation at steady state reflected alterations in recruitment or release rates, we performed FRAP (fluorescence recovery after photobleaching) experiments under the same conditions. Interestingly, the NSRa exchange rate between the nucleoplasm and the Nuclear speckles are similar in cells with or without *ACHLYS* overexpression (Figure S8B). To test if overexpressed *ACHLYS* RNAs accumulate in the nucleus or, if they are exported to the cytoplasm, we fractionated the cytoplasm and nucleus, and quantified RNA levels by qPCR. We could show that in OE-*ACHLYS* lines *ACHLYS* accumulates mainly in the nucleus (92%), similar to our positive controls for nuclear retention SEN1-unspliced (74%) and U6-snRNA (93%). In contrast, our negative control for nuclear retention, SEN1-spliced, is mostly exported to the cytoplasm (10%) (Figure S8C).

Altogether, we have identified several new lncRNAs affecting AS landscapes, notably during LRF and one, *ACHLYS*, was shown to interact with NSRa to regulate specific AS events. Furthermore, our results suggest that overexpression of *ACHLYS* is likely affecting NSRa partitioning into Nuclear speckles *in vivo*, which in turn can affect AS of specific NSRa targets. Hence, *ACHLYS* action defines novel mechanistic insights into the regulatory effects of lncRNA in AS of eukaryotes.

## Discussion

We used a detailed temporal transcriptome of gravitropic stimulus induced LRF and identified hundreds of AS events. While certain of them were detected in genes related to root development, others have no known function in this process. Specific AS patterns define tissue identity in both humans and plants (Baralle and Giudice 2017; Martín et al. 2021) and cell type specific transcriptome and proteome analysis confirmed that AS contributes to root development (P. Lan et al. 2013; Li et al. 2016). Moreover, the characterization of individual AS events proved that specific protein AS isoforms regulate root and lateral root development (Sibout et al. 2006; Remy et al. 2013; Jayaweera et al. 2014). Many time-course RNAseq datasets were generated in the context of plant stress responses (Caldana et al. 2011; Coolen et al. 2016; Calixto et al. 2018) but much less in developmental processes. Reports on the dynamic AS regulation during organogenesis are rare (Filichkin and Mockler 2012; Merens et al. 2024). However, several examples showed that this approach allows the identification of functional important AS events in human development such as time course AS analysis during the organogenesis of brain, heart, liver and kidneys (Mazin et al. 2021). Thus, our genome-wide analysis of AS during lateral root organogenesis also considering lncRNAs contributes to our global understanding of the dynamic regulation of AS during development (Kalsotra and Cooper 2011; Mazin et al. 2021; Lio et al. 2023). Indeed, we identified several AS events induced by lncRNAs that are likely implicated in lateral root development.

In addition to the AS developmental dynamics, we used this dataset to identify several lncRNAs differentially expressed during LRF interacting with the SFs NSRa and GRP7. To maximize the identification of *bona fide* lncRNA regulators of AS we used besides their expression pattern in LRF, additional criteria. We selected monoexonic lncRNAs, to discard the interaction with SF due to intron processing, lncRNAs that interact with the SFs NSRa and GRP7 (Meyer et al. 2017; Bazin et al. 2018), not giving rise to small RNAs based on databases and not being antisense transcripts, which may lead to AS changes through other indirect effects. After transient introduction of these selected lncRNAs in leaf cells, we tested direct effects on AS after 3 days, an early time point where significant expression of the transgene was detected.

This screen led to the identification of *ACHLYS*, a new lncRNA that interacts with NSRa and is involved in AS regulation. Notably, also the known AS regulating lncRNAs *ASCO* and *FLAIL* affected AS in the transient expression assay. The identification of *FLAIL* as an AS-regulating lncRNA independently after our screening by Jin et al. 2023 is a strong support for the effectiveness of our screening strategy. *ASCO* overexpression and *nsra* mutants share similar AS events, suggesting that *ASCO* decoys NSRa from pre-mRNA to modulate their AS (Bardou et al. 2014; Rigo et al. 2020). *FLAIL* in contrast is supposed to recruit NSRa to flowering genes through short sequence motifs, which serve to bind DNA by complementarity and R-loop formation (Jin et al. 2023). When we transiently overexpressed several lncRNAs that interact with the same SFs (NSRa and GRP7), most induced AS are unique for each lncRNA. Only *ACHLYS* and *FLAIL* show partial overlapping on shared AS events, whose dPSI is positively correlated implying a similar way of AS regulation of this subset of genes. In humans, several lncRNAs interact with the same SF. The lncRNAs PANDAR and Pnky both decoy the SF PTBP1, while the lncRNA PNCTR recruits PTBP1 to pre-mRNAs (Pospiech et al. 2018; Yap et al. 2018; Du et al. 2023). Also, the action of the SF SRSF1 is modulated by four lncRNAs affecting its phosphorylation, localization and stability (Tripathi et al. 2010; Ishizuka et al. 2014; H. Wu et al. 2015; Duan et al. 2021). On the other hand, a single lncRNA could also interact with multiple components of the spliceosome. In addition to NSRa, *ASCO* interacts also with core spliceosomal proteins PRP8 and SMD1b and *FLAIL* with U1, U2 and U5 snRNAs (Rigo et al. 2020; Jin et al. 2023). Thus, both of these plant lncRNAs can directly or indirectly bind different core spliceosomal components. In fact, *ACHLYS*, *FLAIL* and *ASCO* interact with NSRa based on NSRa-RIPseq, in which formaldehyde crosslinking can also capture indirect protein-lncRNA interactions (Bazin et al. 2018). The NSRa-iCLIP, using UV crosslinking, allows identification of *in vivo* direct bound RNAs and the location of the NSRa binding sites transcriptome wide. Our NSRa-iCLIP data revealed the direct interaction of NSRa to *ACHLYS* and *FLAIL* through several binding sites. Surprisingly, *ASCO* that was also identified in the NSRa-RIPseq shows no significant NSRa binding sites and only a few NSRa-iCLIP crosslink sites. This could be due to its low expression level in the conditions used for the iCLIP, as iCLIP is biased towards transcript with a higher abundance (Meyer et al. 2017). Alternatively, the interaction detected in RIPseq may be indirect due to formaldehyde crosslinking to other proteins from NSRa ribonucleoprotein complexes. This also occurs with *FLAIL* shown to interact with GRP7 on GRP7-RIPseq, but not on GRP7-iCLIP, supporting indirect interactions of *FLAIL* with GRP7 (Meyer et al. 2017). Interestingly, based on iCLIP data, *ACHLYS* is directly bound by NSRa but also by the SF SR34a, suggesting that also *ACHLYS* could modulate the function of multiple SFs (Laloum et al. 2023). In human medical research, several lncRNAs were also identified to regulate AS through the interaction with the same SF. For instance, SRSF1 phosphorylation status and subcellular localization is regulated by MALAT1 and its stability by the lncRNA DGCR5 (Duan et al. 2021), while the lncRNA Gomafu decoys SRSF1 (Ishizuka et al. 2014) and lincRNA-uc002yug.2 recruits SRSF1 (H. Wu et al. 2015) to target pre-mRNAs. Other examples are the lncRNAs PANDARA and Pnky which both decoy PTBP1 from target pre-mRNA (Pospiech et al. 2018; Du et al. 2023). Furthermore, the SF hnRNPA1 is either decoyed or recruited to target pre-mRNAs by the lncRNAs PCGEM1 and SNHG6, respectively (Z. Zhang et al. 2016; Z. Lan et al. 2020). We could speculate that several lncRNAs interact with the same SF to regulate AS and contribute to define SF selectivity and/or specificity in different biological contexts. AS comparison of transcriptomic data of KD- and OE-*ACHLYS* plants confirmed, in stable plant lines, that ACHYLS regulates AS. Interestingly, the positive correlation between shared AS events in KD- and OE-*ACHLYS* highlights a consistent regulatory effect for these target mRNAs. A similar pattern of shared AS events positively correlated was also observed for *ASCO* knockdown and overexpression (Rigo et al. 2020). Suggesting, that AS regulation by these lncRNAs may rely on a finely tuned stoichiometry with their protein partners. Deviations from this balance, either due to too low or too high lncRNA levels, could disrupt the formation or function of specific lncRNA-protein complexes, ultimately affecting AS fidelity.

To demonstrate that a lncRNA regulates AS through interaction with a SF either individual AS events are characterized in detail (including SF binding to the AS region), or the dPSI of shared splicing events induced by deregulation of the lncRNA or SF are correlated (Romero-Barrios et al. 2018; Marasco and Kornblihtt 2023). To link lncRNAs regulating AS with their interacting SF, we intersected the positional genome-wide information of NSRa RNA binding sites identified by iCLIP with *ACHLYS*- and *FLAIL*-dependent AS events. Our combined approach on lncRNA impact on AS together with NSRa iCLIP, allows us to identify intron retention events whose splicing is presumably regulated by the bound SF (NSRa) as well as the lncRNA. For two *ACHLYS*-dependent intron retention events, we used the transient expression assay to demonstrate that these events could not be detected in *nsra* mutants. This result strongly suggests, at least for these examples, that the AS regulation induced by *ACHLYS* expression depends on *ACHLYS*-NSRa interactions.

The transient expression screen allowed us to address the specificity of the action of lncRNAs. In fact, *ACHLYS* and *FLAIL* were the only lncRNAs that share a substantial part of AS events, which are positively correlated despite that many lncRNAs affect multiple AS events. Similarly, 20% of KD-*ACHLYS* DAS events are also detected in *flail3*, of which are mainly negatively correlated. The differential effect on the common AS events after transient expression or in stable plants could be due also to the different experimental approach (overexpression vs mutations) and different tissues (leaves vs root) that were used. Globally, there seems to be a specific action of lncRNAs in AS despite certain overlapping detected between *FLAIL-* and *ACHLYS*-AS events. For *FLAIL* knockout mutants, nsra and nsra/b an early flowering phenotype was described (Bazin et al. 2018; Jin et al. 2023) and here we could show a root phenotype opposite to the one observed in KD- and OE-*ACHLYS* plants. Thus, *ACHLYS* deregulation and *FLAIL* knockout exhibit also opposite effects on certain shared AS events. We speculate that these AS events contribute to the opposite phenotype. Other lncRNAs are known to be involved in the regulation of root development through other mechanisms than AS regulation. For instance, an antisense lncRNA regulates the expression of Cytokinin biosynthesis gene (Zubko and Meyer, 2007) whereas the lncRNA IPS1 is a miRNA target mimicry sequestering miRNA-399 involved in phosphate homeostasis and root architecture (Franco-Zorrilla et al. 2007). The lncRNA APOLO regulates the expression of its neighboring gene and distal genes epigenetically by the recruitment of chromatin remodelers to target genes by R-loop formation. This mechanism affects root gravitropism and LRF (Ariel et al. 2014; Ariel et al. 2020).

Several SFs are known to undergo LLPS *in vitro* and *in vivo.* It is hypothesized that this property is important for their molecular function as AS regulators (Giudice and Jiang 2024). We showed that the *in vitro* phase separation of NSRa is modulated in a concentration dependent manner by different RNAs including *ACHLYS* - similar to observations in other SFs (Maharana et al. 2018; Wadsworth et al. 2024; Ogura et al. 2025). These *in vitro* results suggest a critical stoichiometry between this lncRNA and NSRa where too much or too low amounts of lncRNA affect LLPS formation paralleling the similar effects observed *in vivo* on the common AS targets of KD- and OE-*ACHLYS* plants. Notably, we could not detect a promoting effect of RNA on LLPS of the NSRa RNA binding mutant, suggesting that direct NSRa-RRM RNA interaction facilitates LLPS and not multivalent electrostatic interactions. Interestingly, different RNA sequences have a similar effect on NSRa LLPS *in vitro*, which is not specific for *ACHLYS*. We could speculate that NSRa interacts with all tested RNAs as the iCLIP deduced binding sites are very small and any RNA contains multiple short degenerated NSRa RNA binding motifs. In contrast, only *ACHLYS* was shown to modulate NSRa accumulation in nuclear biomolecular condensates *in vivo* highlighting a novel potential lncRNA-related mechanism for AS regulation.

Several lncRNAs have been shown to regulate AS by modulating the subnuclear localization of SFs, often through their accumulation in specific nuclear bodies. For instance, the structural lncRNA NEAT1 functions as a central scaffold for protein assemblies of the nuclear biomolecular condensate named paraspeckles (Yamazaki et al. 2018), while MALAT1 regulates the distribution of SR proteins between Nuclear speckles, nucleoplasm and cytoplasm (Tripathi et al. 2010; Miyashita et al. 2022). Motivated by these precedents, we investigated whether *ACHLYS* might influence the subnuclear localization of its interacting partner NSRa. We found that *ACHLYS* overexpression leads to increased accumulation of NSRa in Nuclear speckles, suggesting a possible mechanism by which *ACHLYS* could modulate AS regulation through spatial reorganization of SFs. Notably, *ACHLYS* also accumulates in the nucleus under overexpression conditions, raising the possibility that the lncRNA itself may contribute to NSRa speckle enrichment, either directly through recruitment or indirectly by altering NSRa binding dynamics or nuclear trafficking. Despite the observed steady-state accumulation, FRAP analysis revealed no major differences in the exchange dynamics of NSRa in speckles upon *ACHLYS* overexpression. This suggests that the increase in speckle-associated NSRa is not due to changes in its binding kinetics but may instead reflect a shift in the equilibrium distribution, potentially caused by increased nuclear retention or sequestration *via* stable interactions promoted by *ACHLYS*. Overall, these findings support a model in which *ACHLYS* regulates AS not only through biochemical interactions with NSRa but also by influencing its subnuclear localization.

We propose that *ACHLYS* is a new lncRNA that regulates AS through direct interaction with NSRa and/or by modulating their spatial organization in Nuclear speckles *in vivo*, and revealing the complex network of lncRNAs and SF controlling AS. These results open wide perspectives for the potential of lncRNAs to regulate AS events in eukaryotes.

## Materials and methods

### Plant related materials and methods

#### Growth conditions

If not otherwise indicated, all plants were grown as follows: Wild type (Columbia-0) and transgenic seeds were gas sterilized and vernalized (2-3 days at 4°C). Seeds were placed on plates containing ½ Murashige and Skoog (MS) medium including vitamins (Duchefa), 1.2 % w/v agar (pH 5.6) and moved to a Aralab climate chamber with long day condition (16 h light/ 8 h dark, 20°C, 60% humidity). We refer to this timepoint when we describe seedlings and plants’ age. The plant lines used are summarized in Table S3.

#### Transient expression

The transient expression in wild type (Columbia-0) or nsra *A. thaliana* leaves was performed according to Y. Zhang et al. 2020. Agrobacterium (GV3101 or AGL1) clones containing a vector to overexpress the lncRNA candidates, *ASCO* or GFP as control, were grown in liquid culture overnight (28°C). 200 µl of liquid culture was transferred to YEB-induced medium plates (0.2 mM acetosyringone) and incubated at 28°C for 36 h. Bacteria were scraped from the plate and resuspended in 2 ml washing solution (10 mM MgCl_2_, 100 μM acetosyringone) and diluted in infiltration solution (¼MS [pH = 6.0], 1% sucrose, 100 μM acetosyringone, 0.005%_v/v_ Silwet L-77) to an OD600 of 0.5. The previously marked seventh leaf of 30-35-day-old plants (long day, day 20°C, night 18°C, 60% humidity) was infiltrated with agrobacteria using a 1-ml plastic syringe. The infiltrated plants were dried for 1h in the growth cabinet, incubated in the dark for 24 h and the samples were collected 88 h after infiltration.

#### Root phenotyping

Seedlings were grown for 10 and 14 days after transfer to light, plates were scanned and main root lengths and lateral root lengths were measured using RootNav 2 (Yasrab et al. 2019). Lateral root length is the sum of the length of all lateral roots in one plant and total root system length is the sum of main root length and lateral root length. Lateral root density was calculated by dividing the lateral root number by the main root length.

### Molecular biological methods

#### Cloning

All lncRNA candidates and controls were amplified (using primers summarized in Table S4) and cloned using the GreenGate system (Lampropoulos et al. 2013) into entry vector pGGC000 and combined with pGGA004 (35S Cauliflower Mosaic Virus (CaMV) promoter), pGGB003 (dummy), pGGD002 (dummy), pGGE009 (ubiquitin10 terminator), pGGF007 (pNOS:KanR:tNOS) and pGGZ001 (destination vector). To generate RNAi lines, a sense and antisense fragment of 368 bp at position + 43-311 bp relative to the *ACHLYS* TSS was cloned in pGGB000 and pGGD000, respectively. These fragments were combined with pGGA004 (35S promoter), an inhouse pGGC (containing intron of chalcone isomerase gene, based on pFRN binary destination vector) pGGE009 (ubiquitin10 terminator), pGGF007 (pNOS:KanR:tNOS) and pGGZ001 (destination vector).

#### Gene expression and splicing analysis by qPCR

If not further specified, we used pools of 10-day-old-seedlings for all qPCR analysis. Following the manufacturer’s instructions total RNA was extracted using TRI REAGENT® (Molecular Research Center, Inc.), RNA was treated with DNase I (Thermo Scientific) or TURBO™ DNase (Life Technologies) and reverse transcribed by Maxima H Minus Reverse Transcriptase (Thermo Scientific) using Oligo (dT) primers or a 1:1 mix of Oligo (dT) primers and Random Hexamer Primer. The used *FLAIL* qPCR primers bind outside of the characterized *FLAIL* antisense transcript. qPCR was performed using LightCycler® 480 SYBR Green I Master reagent following the manufacturer’s instructions using gene specific primers and LightCycler® 96 (ROCHE) or LightCycler® 480 II (ROCHE) following standard protocol. The efficiency of all primer pairs was tested and DNase I (Thermo Scientific) treated but not reverse transcribed RNA samples were used to detect genomic DNA contaminations. The number of biological replicates is indicated for each experiment ranging from 3 to 8.

For qPCR analysis the ΔC_t_ was calculated as C_t_ reference gene (PP2A, AT1G13320) minus C_t_ gene of interest for each biological replicate. Higher ΔC_t_ values indicate a higher expression of the gene of interest compared to the control. To test statistically significant differences between ΔC_t_ values of different samples we used an unpaired, two-tailed T-test (*e.g.*, mean ΔC_t_ transgenic compared to mean ΔC_t_ control). Direct expression differences between mutant and wild type are shown as ΔΔC_t_ (mean ΔC_t_ transgenic – mean ΔC_t_ control) and standard error of mean difference, which were obtained from the statistical analysis. Used primers are summarized in Table S5.

For splicing qPCR, we used three primer pairs per gene: First amplifying a shared exon, as an internal normalizer; second, using primers spanning the exon–exon junction to quantify the spliced mRNA; and third, using primers located within the retained intron to detect intron retention (IR) events. To calculate the ΔC_t_ for different RNA isoforms, we calculated C_t_ of shared exon minus C_t_ spliced or C_t_ retained intron for each biological replicate and compared the mean difference between biological replicates of the control to different samples (*e.g.*, ΔΔC_t_ = mean ΔC_t_ transgenic – mean ΔC_t_ control. Mean difference shows ΔΔC_t_ and standard error of mean difference, which were obtained from the statistical analysis. Used primers are summarized in Table S6,7.

#### Library preparation for transient expression screen and KD- and OE-*ACHLYS*

Total RNA was extracted from individual seventh leaves of 37-day-old-plants grown in an Aralab climate chamber with long day conditions (16 h light/ 8 h dark, 20°C, 60% humidity), or from pools of 10-day-old seedlings for KD- and OE-*ACHLYS*. TURBO™ DNase (Life Technologies) treated RNA was purified using the RNA Cleanup protocol of the RNeasy Plant Mini Kit (QIAGEN) and used for Illumina® Stranded mRNA Prep (Illumina) following the manufacturer’s instructions. We performed a second library clean up to reduce primer dimer contamination. Libraries were sequenced by BGI genomics (China)

#### NSRa-iCLIP

The NSRa-iCLIP was performed according to plant iCLIP2 protocol (Lewinski et al. 2024). Plants expressing the NSRa-GFP fusion protein under control of the endogenous promoter in the nsra mutant background were grown for 14 days in 12 h light-12 h dark cycles. Seedlings were shifted to 38°C for 3 hours and subjected to crosslinking with 254 nm UV light at 4000 mJ/cm^2^. Whole seedlings were harvested, flash frozen and ground to a fine powder. RNA-protein complexes were immunoprecipitated from crude lysates with GFP Trap beads (Proteintec, Martinsried, Germany). An aliquot of the precipitate was treated with RNase on the beads. The RNA was radioactively labeled. RNA-protein complexes of both the samples and the RNase control were separated by denaturing gel electrophoresis, transferred to a Nitrocellulose membrane and subjected to autoradiography. Disappearance of the radioactive smear above the expected size for the fusion protein upon RNase treatment demonstrated that RNA has been crosslinked. The region above the expected size for the fusion protein was cut out and subjected to proteinase K digestion. RNA was isolated for library preparation. The first iCLIP replicate was performed with 20 g yielding 2.7 Mio deduplicated read. For the second and third replicate we scaled up the procedure and performed iCLIP on 70 g, each resulting in up to 21 Mio reads.

#### RNA Electrophoretic Mobility Shift Assay (RNA-EMSA)

NSRa-RRM a was cloned in the bacterial expression vector pGEX-6P1 (GE Healthcare, Freiburg, Germany) and GST-NSRa-RRM-(FD) mutant was generated by site-directed-mutagenesis of pGEX-6P*1-NSRa* vector using complementary mutagenic primers according to Zheng et al. 2004. Recombinant GST-tagged NSRa-RRM and NSRa-RRM-(FD) were purified from BL21 cells after IPTG induction (0.2 mM) of cell cultures with OD600 of 0.5 and 3 h of incubation at 30°C, using Pierce™ GST Spin Purification Kit following the manufacturer’s instructions. For the RNA EMSA assay proteins were incubated with 3′-biotinylated RNA probes (22 nucleotides) corresponding to a sequence within the main NSRa iCLIP peak on the lncRNA *ACHLYS* (UGCCGCCGCUUAGUGAAAAAUU). A scrambled RNA probe with identical nucleotide composition (GGUGCUUGCGUAACAUCAUACA) was used as a control. Binding reactions were carried out in a final volume containing 10 mM HEPES (pH 7.3), 20 mM KCl, 1 mM MgCl₂, 1 mM DTT, 5% glycerol, 2 µg tRNA, 6.25 µM recombinant protein, and 25 nM biotin-labeled RNA probe. Reactions were incubated for 30 min at room temperature and subsequently resolved on native 6% polyacrylamide gels prepared in 0.5× TBE buffer. Electrophoresis was performed at 100 V for 40 min, followed by transferring to a nylon membrane for 35 min at 400 mA. RNA–protein complexes were detected using the LightShift® Chemiluminescent RNA EMSA Kit (Thermo Fisher Scientific) according to the manufacturer’s instructions, using streptavidin–horseradish peroxidase conjugate for signal development.

#### *In vitro* NSRa phase separation assay

Phase separation assays were performed using recombinant NSRa proteins purified as described above. To induce *in vitro* droplet formation, we mix 6 µM purified GST-NSRa-RRM, or GST-NSRa-RRM-(FD) with 100 mM KCL, 10% PEG (8000), 15 mM NaCl, 5 mM TRIS base (pH 8.0), 2 % glycerol and 1 mM Reduced Glutathione. RNAs were *in vitro* transcribed, using HiScribe® T7 High Yield RNA Synthesis Kit, from templates generated using a PCR product of the gene of interest including T7 promoter sequence encoded in the forward primer (Table S8).

#### Microscopy-based analysis of NSRa subnuclear localization

For the analysis of NSRa subnuclear localization, we used *A. thaliana* seedlings carrying a genomic NSRa-GFP fusion expressed under its native promoter in a nsra knockout background. Seedlings were grown for 6 days under long-day conditions, and epidermis cells of the main root meristem were analyzed. To assess the effect of lncRNA overexpression, we generated stable transgenic lines by transforming nsra pNSRa:NSRa-GFP plants with constructs that constitutively express the lncRNAs of interest under the 35S promoter. Confocal imaging (5 µm of pinhole) was performed on root tips using the 488nm laser (2% intensity) of a Zeiss microscope model LSM880 with a 63x objective. Nuclear speckles were identified based on the punctate GFP-NSRa pattern, previously shown to co-localize with the speckle marker SR34 (Bardou et al. 2014). Image analysis was performed using CellProfiler (version 4.2.6). A custom pipeline was developed to automatically segment speckles and nucleoplasmic regions based on fluorescence intensity. Segmentation accuracy was visually validated by comparing outlines generated by the software with raw images (see Figure 6). For each cell, mean fluorescence intensity was calculated separately for speckles and nucleoplasm, and speckle enrichment was expressed as the ratio of these two values. Between 30 and 60 cells from 5 to 10 independent seedlings were analyzed per condition.

#### Fluorescence Recovery After Photobleaching (FRAP)

FRAP experiments were performed on epidermis cells of the main root meristem of nsra pNSRa:NSRa-GFP and nsra pNSRa:NSRa-GFP OE-ACHYLS plants to evaluate the dynamic exchange of NSRa in Nuclear speckles. Confocal imaging was conducted using the 488 nm laser (2% intensity) of a Zeiss microscope model LSM880 with a 63x objective. A single Nuclear speckle per cell was photobleached using the 488nm laser at 12% of intensity and fluorescence recovery was recorded at 459 msec frame rate for a total time of 5 sec. At each time point, speckle fluorescence intensity was normalized to a non-bleached nucleoplasmic region to correct for acquisition photobleaching. Pre-bleach intensity was calculated as the average of three frames acquired before bleaching, and recovery was expressed as the percentage of this initial value over time. Average recovery curves were generated from at least 40 speckles per condition, each from an independent cell.

### Bioinformatic methods

#### Bioinformatic screen for lncRNA candidates with potential function in alternative splicing

The source data for the bioinformatic screen for lncRNAs with potential function in regulatory AS networks are two datasets: First RIPseq data for NSRa (Bazin et al. 2018) and iCLIP data for GRP7 (Meyer et al. 2017). All NSRa and GRP7 interacting lncRNAs that have in the TAIR10 reference genome an associated antisense transcript, a major isoform that is spliced or precursors for small RNAs, were excluded. Five candidates interacting with NSRa and six candidates interacting with GRP7 were individually selected based on their differential expression during LRF. Two additional lncRNA which were not detected in the GRP7-iCLIP and NSRa-RIPseq were selected.

#### Transcriptome analysis

Libraries were prepared using a Tru-Seq Stranded mRNA Sample Prep kit (Illumina®). Sequencing was performed using a paired-end protocol to a minimum depth of 40 million reads per sample on an DNB-seq 100nt paired end (BGI Genomics, China). Raw reads were quality-trimmed and adapter sequences removed using **Cutadapt** (5.1). Cleaned reads were aligned to the *A. thaliana* reference genome using **STAR** (2.7.11b), with minimum and maximum intron lengths set to 50 and 5000 nt, respectively. Only uniquely mapping reads were retained for downstream analysis. Gene-level expression quantification was performed using **HTSeq** (2.0.5) in union mode, using the **Araport11** gene annotation. Differential gene expression analysis was carried out using **DESeq2**, with pairwise comparisons between genotypes. Genes were considered differentially expressed if they had a false discovery rate (FDR) < 0.01.

#### Transcript Quantification and Differential Alternative Splicing Analysis

Transcript-level abundance estimation was performed using **Kallisto** with 100 bootstrap. Quantification was based on the **AtRTD2** reference transcriptome (R. Zhang et al. 2017). Differential alternative splicing (DAS) analysis was conducted using **SUPPA2** (Trincado et al. 2018), using transcript per million (TPM) values from Kallisto. Splicing events were considered differentially regulated if they exhibited an absolute dPSI (|dPSI|) > 0.2 and FDR < 0.01.

To identify isoform switching events in the **lateral root time series**, transcript-level TPM values from **Kallisto** were analyzed using the **TSIS** R package (W. Guo et al. 2017). TSIS was used to detect transcripts that switched their relative expression dominance over time, indicating isoform switching during lateral root development.

#### Characterization of *ACHLYS*

To identify *ACHLYS* homologous in other plant species, we used the NCBI nucleotide Blast and queried *ACHLYS*A against the Reference RNA sequences of Braccicasea using discontiguous megablast using default settings. The retrieved

#### Gene ontology analysis

Gene ontology analysis was performed with indicated gene list using ShinyGO 0.82 and default settings.

#### Bioinformatics analysis of iCLIP

Sequencing reads were demultiplexed and underwent the first quality control (QC). The quality control was performed by using **FastQC** (0.11.9), and the distribution of sample barcodes was computed using **AWK** (1.3.4). The adapter removal and demultiplexing of reads was performed using **Flexbar** (3.5.0), as well as discarding reads shorter than 15 nt. Afterwards, a second QC step was performed with **Flexbar** (3.5.0) as well as a further quality trim (removing reads shorter than 15 nt). A third QC with **FastQC** (0.11.9) was then performed to confirm the quality of the reads for genome mapping. The unique molecular identifiers (UMIs) were added to the fastq file by using **BioAWK** (20110810). Then, a genome index was created by using **STAR** (2.7.3a) using the *A. thaliana* TAIR10 genome and the **AtRTD3** gene annotation (Zhang et al. 2017). The mapping itself was done using **STAR** (2.7.3a). The resulting BAM files were then indexed using **samtools** (1.13.4), and the PCR duplicates were removed based on the UMIs with **umi_tools** (1.1.4). To generate the crosslink tracks to visualize the data, the files were processed using **AWK** (1.3.4), **bedtools** (2.30.0) and **bedGraphToBigWig**. Afterwards, peak calling was performed using the mapped alignment (BAM) files. First off, deduplicated reads were merged and the resulting file indexed by using **samtools** (1.13.4). Then, peaks were called from the uniquely mapped and deduplicated reads using **PureCLIP** (1.3.1) (Krakau et al. 2017).

Then, binding sites were defined based on the called peaks together with crosslink signals from each individual replicate. This was accomplished by using the R-package **BindingSiteFinder** (1.7.2), as well as the current versions of the packages rtracklayer, GenomicFeatures, tidyr and dplyr. An initial BindingSiteFinder object was created using this data, with meta-information provided for each sample. To reduce noise, peaks with the lowest 1% PureCLIP scores were globally removed using the **pureClipGlobalFilter** function. Gene annotations from AtRTD3 and Ensembl were then imported and linked via unique gene IDs. Using the AtRTD3 annotation, a TxDB database was generated, allowing for the retrieval of gene and transcript regions. Positions for all annotated genes were extracted.

The estimated BsWidth function was employed to simultaneously determine parameters for binding site size (bsSize) and gene-wise filtering (bsFilter). A gene-wise filter was then applied, retaining the top 50% of PureCLIP peaks per gene while discarding the lowest 50%.

A set of equally sized binding sitesof 3 nucleotides width was computed using the function makeBindingSites. To account for variability among replicates, a replicate reproducibility filter was then applied. Binding sites had to pass the 10% quantile threshold of crosslinks per binding sites for each replicate in at least 2 out of 3 cases.

Binding sites were assigned to their respective host genes based on overlaps. Specific transcript regions including coding sequences (CDS), introns, and untranslated regions (UTRs) were extracted from the annotation database. Binding sites were then mapped to their corresponding transcript regions. Each binding site was reassigned a PureCLIP score by selecting the highest score from all overlapping PureCLIP peaks. Finally, a BED file containing transcriptome-wide binding sites for the processed sample was exported.

#### Enrichment for iCLIP binding site

To quantify the NSRa-iCLIP binding sites around the splice site of lncRNA dependent intron retention events we used an in-house R script. We quantified the presence of an NSRa-iCLIP peak in a window 50 nt downstream and 50 nt upstream of the 5’- and 3’-SS of a lncRNA dependent intron retention events identified by **SUPPA2** (Trincado et al. 2018). As a control we quantified the NSRa-iCLIP binding site in all annotated intron retention events of the **AtRTD3** gene annotation (Zhang et al. 2017).

#### General usage of statistics

All statistics and graphs were created using R or Prism8 (GraphPad Software). Unpaired two-tailed student T-test was performed to compare two samples. One-way analysis of variance (ANOVA) and Tukeýs honestly significant difference (HSD) post-hoc test was used to compare more than two samples.

## Supporting information

Supplementary_results

## Data availability

The RNAseq of the transient expression screen, KD- and OE-*ACHLYS* lines and corresponding controls have been deposited in GEO (http://www.ncbi.nlm.nih.gov/geo) with accession number GSE318723. The three NSRa-iCLIP replicates have been deposited in the Sequence Read Archive (SRA) under the accession numbers SRR36997292, SRR36997293, and SRR36997294.

## Acknowledgment

This work was supported by the ANR-DFG bilateral RIBORES grant (ANR-22-CE92-0018 to M.C. and STA653/18-1 to DS) and an FRM fellowship (ECO202106013730/FRM) to MH. SM acknowledges the funding from the Novo Nordisk Foundation (NNF19OC0057485, NNF21OC0066776). The MC Lab at IPS2 benefits from the support of Saclay Plant Sciences-SPS (ANR-17-EUR-0007).

## Author contributions

MH, TB, PM and AC performed and analyzed experiments. ML, JB, TB performed bioinformatic analyses. JB, DS and MC designed the experiments; MH, JB, DS MC conceived the study and wrote the manuscript. MC and DS acquired funding.

## Supplementary figures

**Figure S1:**
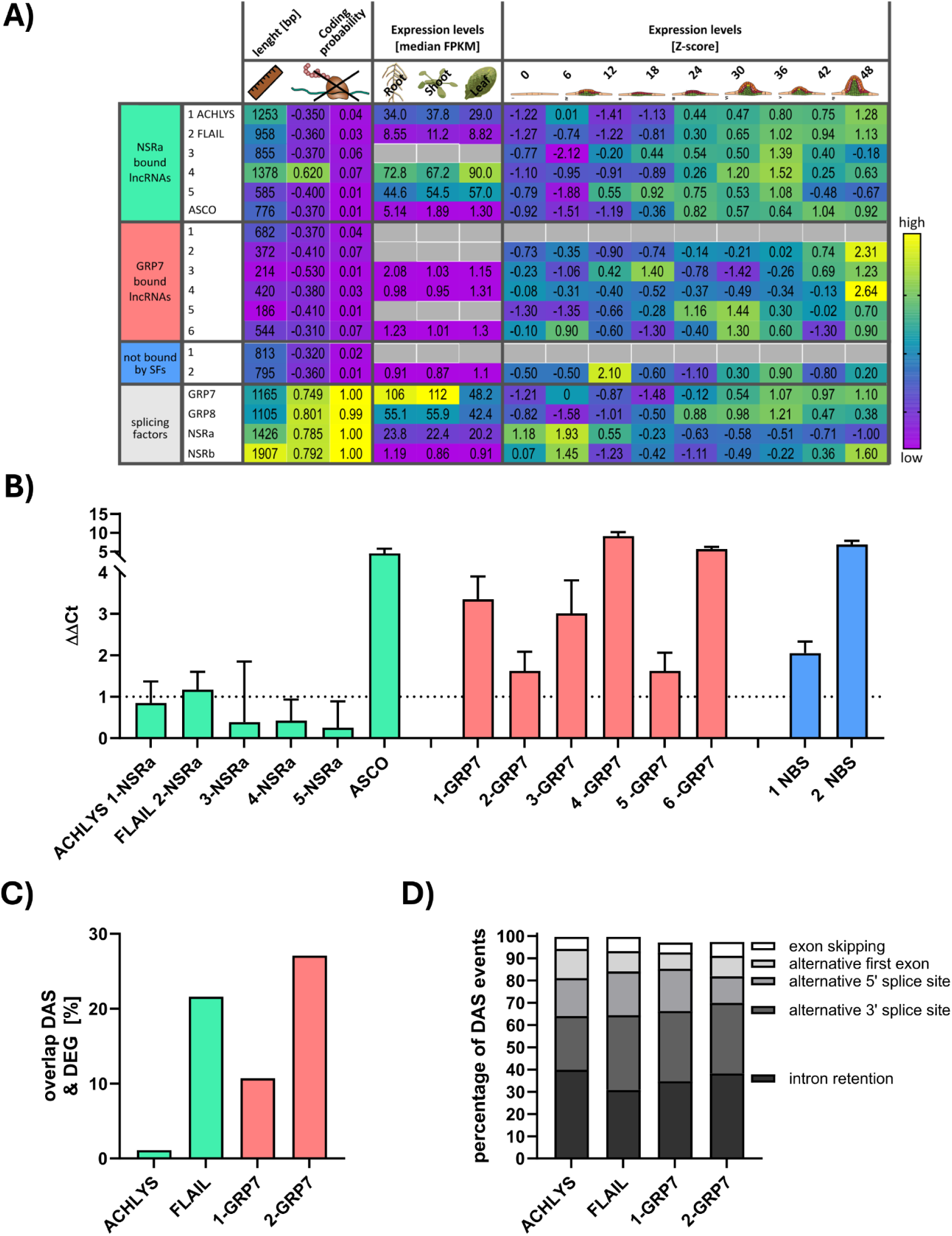
Transient expression screen. **A)** Characterization of lncRNA candidates. Table displays lengths, coding probability calculated according to the CNIT (left) and CPC2 (right). Median FPKM (fragments per kilobase of transcript per million mapped reads) levels in root, shoot and leaves. Expression levels, shown as Z score generated from transcript per million (TPM) values of characterized genes during lateral root formation 0-48 hours after induction by root bending. **B)** RNA levels represented as ΔΔCt after transient overexpression in seventh leaf of lncRNA candidates normalized to their expression in plants transiently overexpressing GFP. **C)** Percentage of differentially alternatively spliced (DAS) genes being also differentially expressed genes (DEG). **D)** Percentage of alternative splicing events per type of event detected after transient expression of *ACHLYS*, *FLAIL*, *1-GRP7*and *2-GRP7*. The splicing event alternative first exon is not depicted.

**Figure S2:**
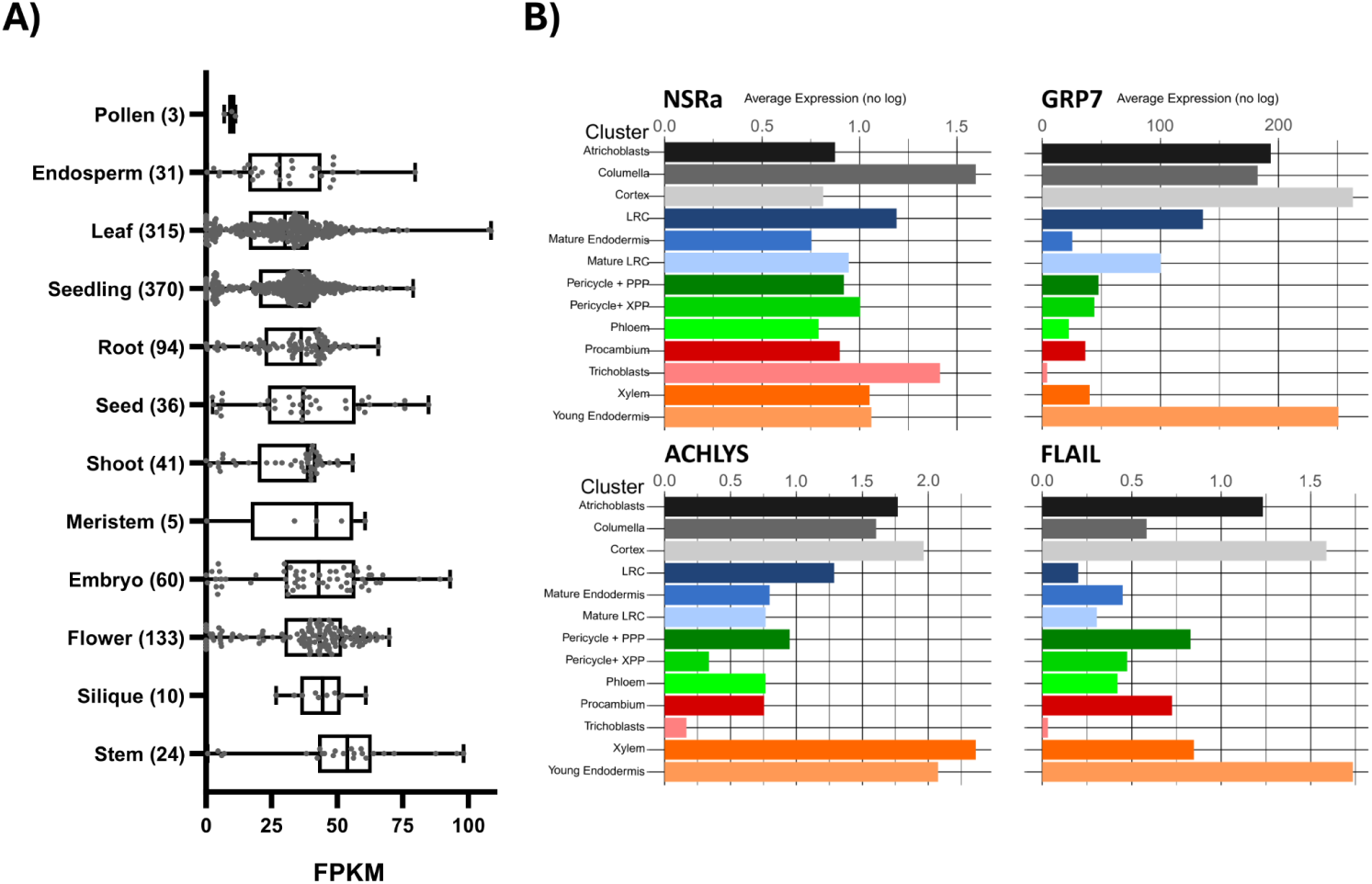
*ACHLYS* expression levels in different organs and root cell types. **A)** *ACHLYS* expression levels as FPKM (fragments per kilobase of transcript per million mapped reads) in different organs and tissues based on Arabidopsis RNA-seq Database (H. Zhang et al. 2020). Numbers of used RNAseq datasets are shown in brackets. **B)** Average expression levels of *ACHLYS*, *FLAIL*, NSRa and GRP7 in different root cell types, determined by single cell RNAseq from (Wendrich et al. 2020).

**Figure S3:**
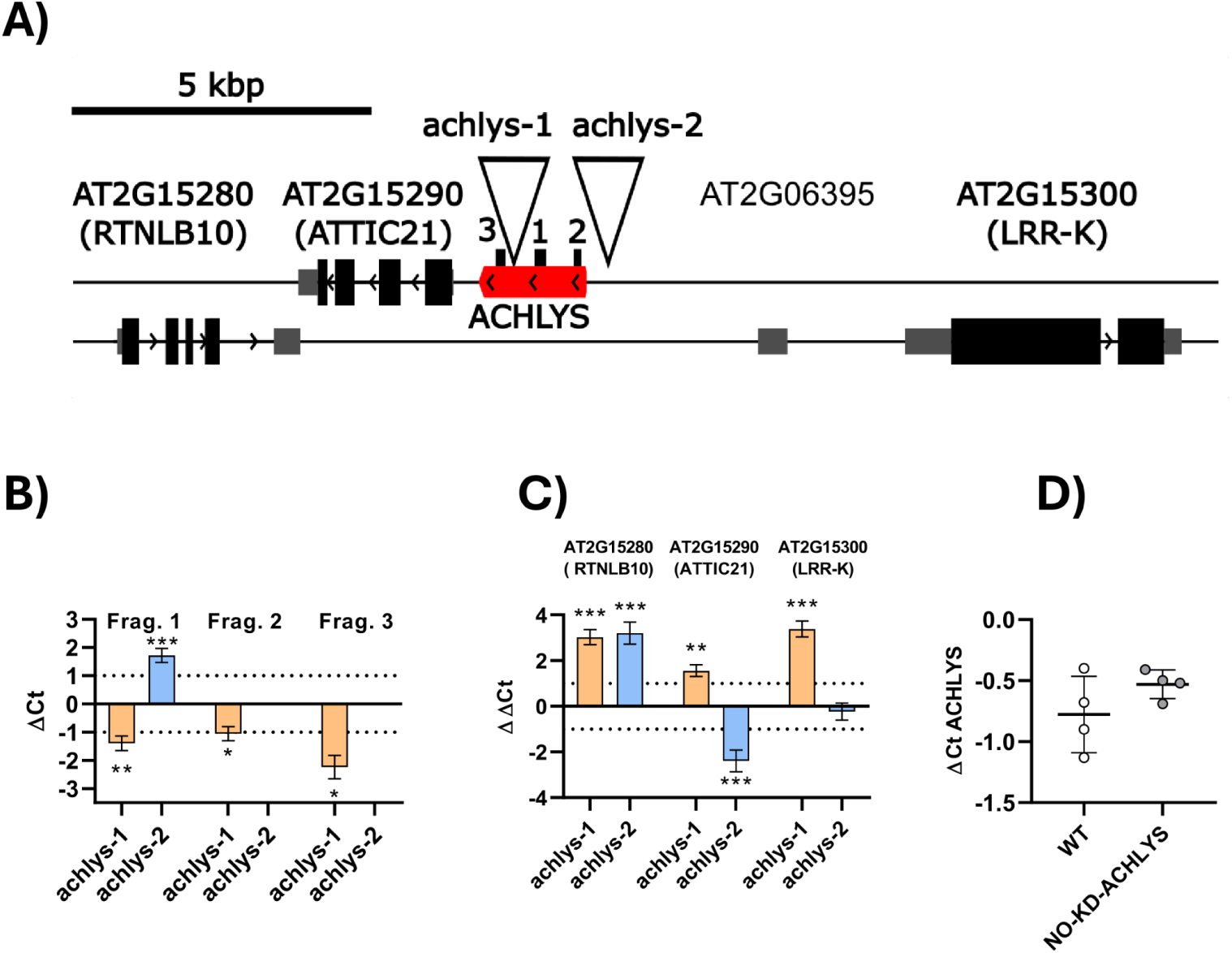
RNA levels of *ACHLYS* neighboring genes in *ACHLYS* plants carrying T-DNA insertions around this locus. **A)** *ACHLYS* locus (red) and neighboring genes. T-DNA insertion sites are shown as white triangles and PCR fragment positions are shown as black lines. Exons are shown as bars (URTs, grey and CDS, black) and introns as lines. **B)** *ACHLYS* RNA levels in T-DNA lines using different primer pairs in 10-day-old seedlings. **C)** qPCR results showing ΔΔCt for *ACHLYS* neighboring genes detected compared to corresponding wild type plants in l *ACHLYS*-1 and -2 showing the impact of the T-DNA insertion on neighboring genes. Dotted lines indicate two-fold change. ΔΔCt as mean difference (ΔCt mutant minus ΔCt wild type comparing 3 to 4 biological replicates and standard error of mean (unpaired, two-tailed T-test, *p ≤ 0.05, **p ≤ 0.01, ***p ≤ 0.001). **D)** qPCR results showing wild type *ACHLYS* levels in 35S:RNAi-*ACHLYS* line non-KD-*ACHLYS* as ΔCt.

**Figure S4:**
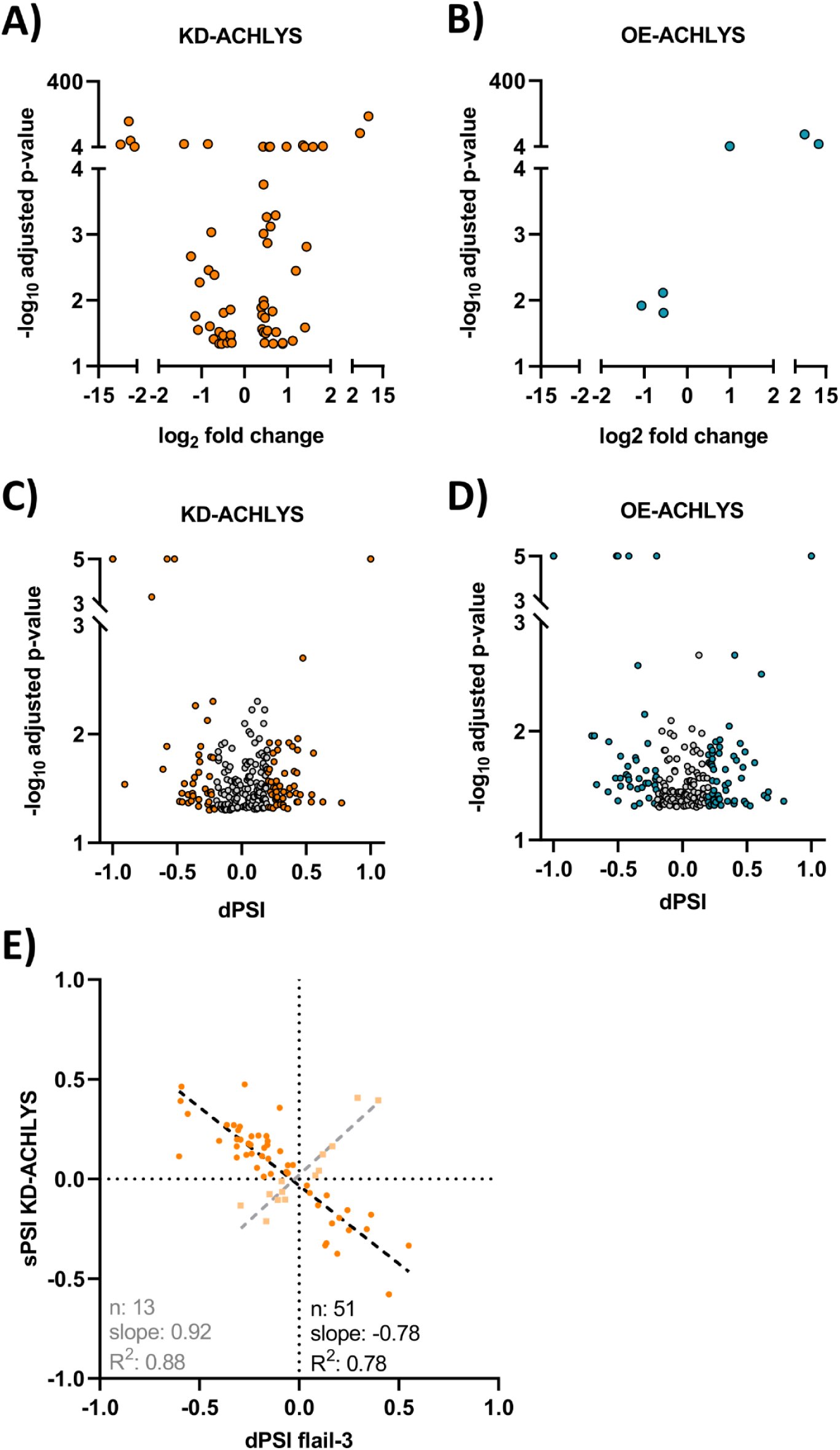
Effect of *ACHLYS* deregulation on gene expression and alternative splicing. **A)** Volcano plot showing log_2_ fold changes in gene expression and -log_10_ adjusted p-value for individual genes (dots) in KD and **B)** OE-*ACHLYS*. **C)** Volcano plot showing delta percent spliced in (dPSI) and -log_10_ adjusted p-value quantified by SUPPA2 in KD-and **D)** OE-*ACHLYS*. Each dot represents one alternative splicing event. Grey dots indicate splicing events with dPSi between 0.2 and - 0.2. **E)** Correlation between dPSI of shared AS events detected in *flail3* mutant and KD-*ACHLYS*. Number of DAS events (n), slope and R2 of the linear regression are shown.

**Figure S5:**
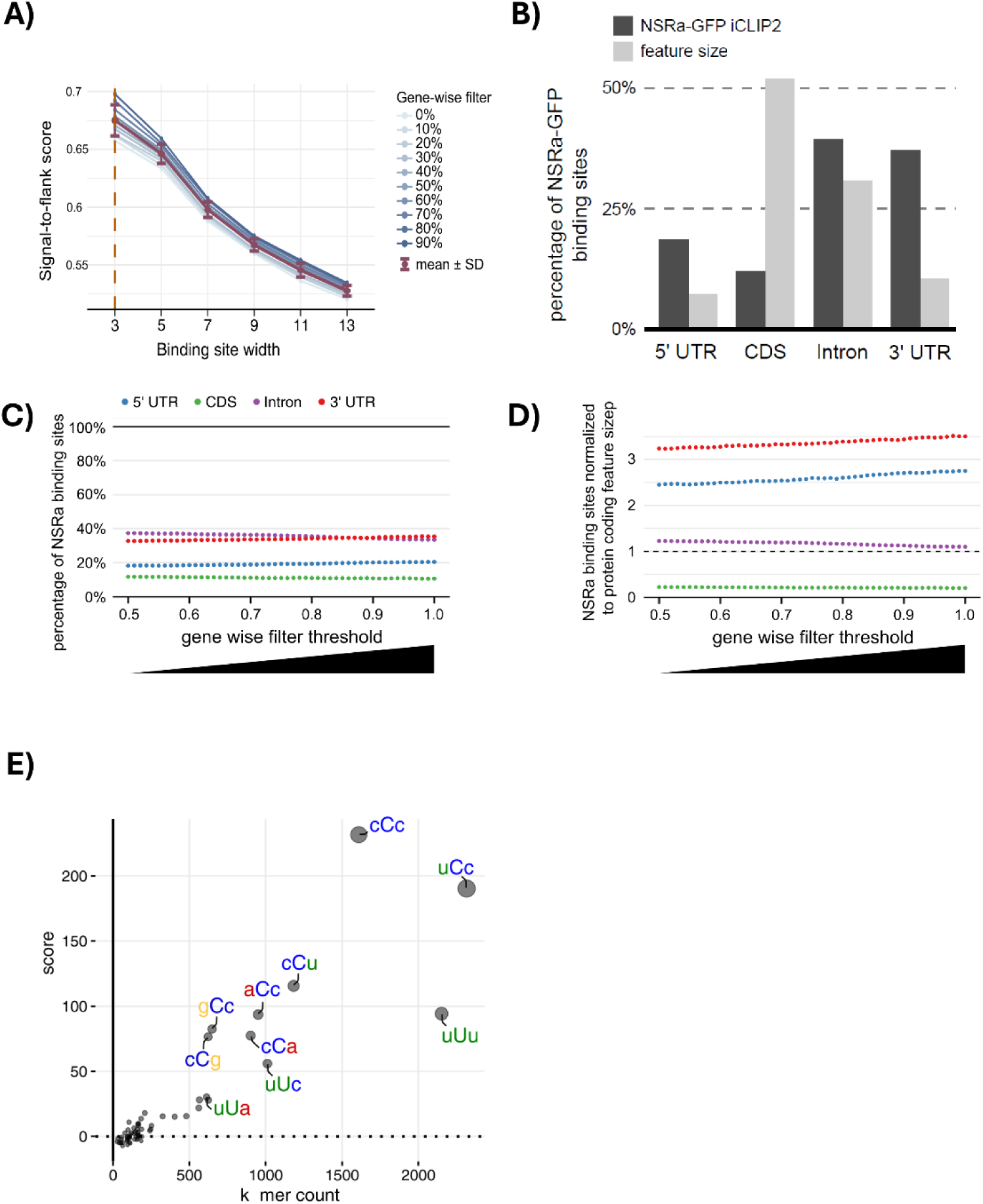
Distribution of NSRa-GFP binding sites within protein-coding transcripts and binding site width. **A)** Empirical estimation of binding site width after increasing gene-wise filter cutoffs (light to dark blue), with y-axis depicting the signal-to-flank score and x-axis describing growing binding site widths (3 to 13 nucleotides). Mean and standard deviation values for each width are shown in brown. The orange dashed line highlights the chosen binding site width for NSRa-GFP. **B)** Percentage of NSRa-iCLIP binding sites in different protein coding transcript features (dark grey bars) compared to the percentage of cumulative length of the indicated region (light grey bars) based on Araport11 gene models. **C)** Distribution of binding sites in different transcript regions (dotted lines) in relation to the score of the binding site peak. The colored dots correspond to reproducible binding sites mapping to the 5’ UTR (blue), coding region (green), introns (purple), and 3’-UTR (red), respectively. The x-axis describes the progressing gene-wise filtering threshold, i.e., how the distribution within the transcripts forms after removing the lowest scored binding sites in comparison to the highest measured peak-score of a transcript. The y-axis shows the percentage of NSRa-GFP binding sites matching the filtering threshold of the x-axis. **D)** Enrichment of reproducible binding site distribution normalized by the cumulative length (feature size) of the indicated region. The dashed line marks an expected distribution according to the cumulative length of the region. **E)** Sequence motifs enriched at the NSRa-iCLIP binding sites. Scatterplot displaying k-mer counts (x-axis) and k-mer z-scores (y-axis). K-mers in the upper right corner are enriched in comparison to a random background correction model. Letters in upper case mark the peak position of the reproducible binding site.

**Figure S6:**
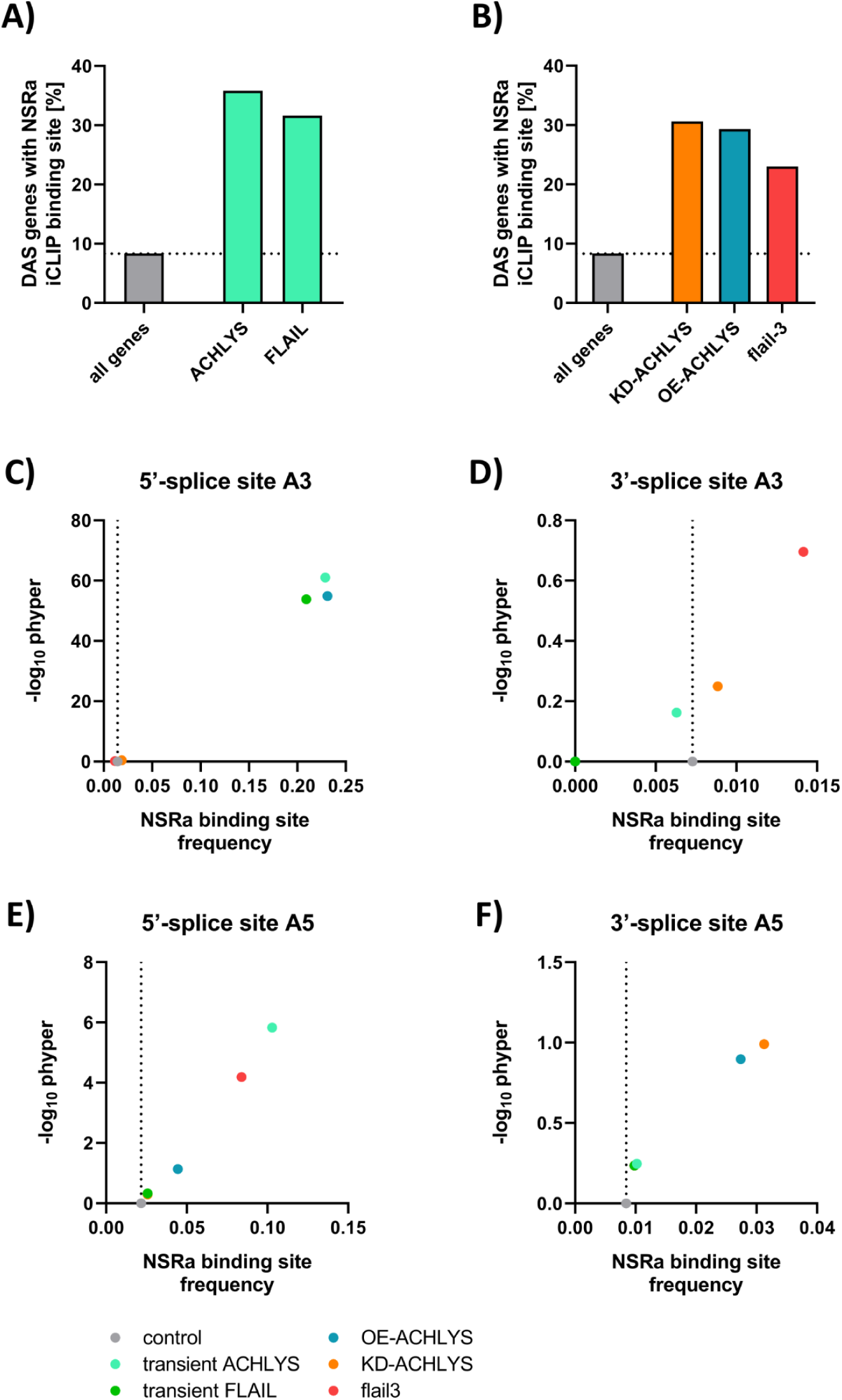
Percentage of ACHYLS and *FLAIL* dependent AS genes with NSRa binding sites. **A)** Percentage of ACHYLS and *FLAIL* dependent AS genes induced by transient ACHYLS and *FLAIL* overexpression with NSRa binding sites. **B)** Percentage of ACHYLS and *FLAIL* dependent AS genes with NSRa binding sites for stable transgenic lines. The gray bar represents the percentage of genes with NSRa binding sites in relation to all genes. **C-F)** X-axis showing NSRa-iCLIP binding site frequency in a 100 nt window centralized at the 5’- and 3’-splice sites of different alternative splicing events in leaves transiently overexpressing *ACHLYS* and *FLAIL* or lines with deregulated *ACHLYS* and *FLAIL* levels. Statistical significance was tested by Hypergeometric test and -log_10_ phyper values are displayed as Y-axis. As a control we used the NSRa binding site frequency at all annotated alternative 5’-(A5) and 3’-splice site (A3) events (dotted line).

**Figure S7:**
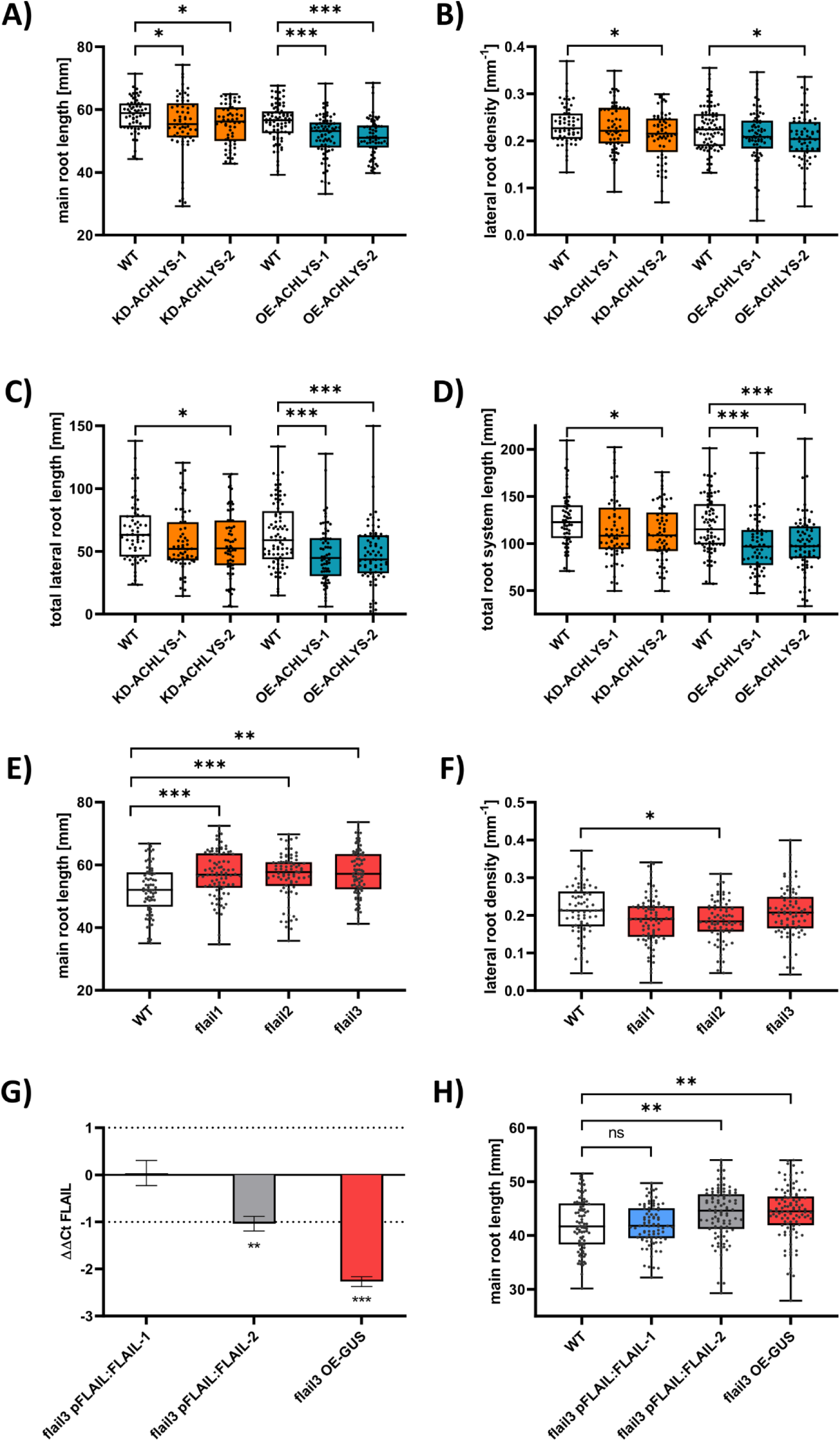
Correlation between *ACHLYS* and *FLAIL* level and root development. **A)** Main root length, **B)** lateral root density, **C)** Total lateral root length and **D)** total root system lengths as sum of main root length and total lateral root length in 14-day-old seedlings of lines with deregulated *ACHLYS* level. Compiled data of three independent experiments (n = 61-81). **E)** Main root length and **F)** lateral root density in 14-day-old seedlings in *FLAIL* knockout lines (CRISPR deletion *flail1* and *flail2* and T-DNA insertion mutant *flail3*) Compiled data of three independent experiments (n = 78-92). **G)** *FLAIL* RNA levels in 10-day-old seedlings in *flail3* complementation lines and GUS control. ΔΔCt as mean difference (ΔCt mutant minus ΔCt control, comparing 3 to 4 biological replicates and standard error of mean difference, Dotted lines indicate two-fold change (unpaired, two-tailed T-test, *p ≤ 0.05, **p ≤ 0.01, ***p ≤ 0.001). **H)** Main root length in 10-day-old seedlings in *flail3* complementation lines and controls. Compiled data of three independent experiments (n = 88-99). Box-and-whisker plots (median, first and third percentiles, whiskers min-max percentile, dots represent biological replicates), one-way analysis of variance (ANOVA) and Tukeýs honestly significant difference (HSD) post-hoc test, *p ≤ 0.05, ** p ≤ 0.01., ***p ≤ 0.001.

**Figure S8:**
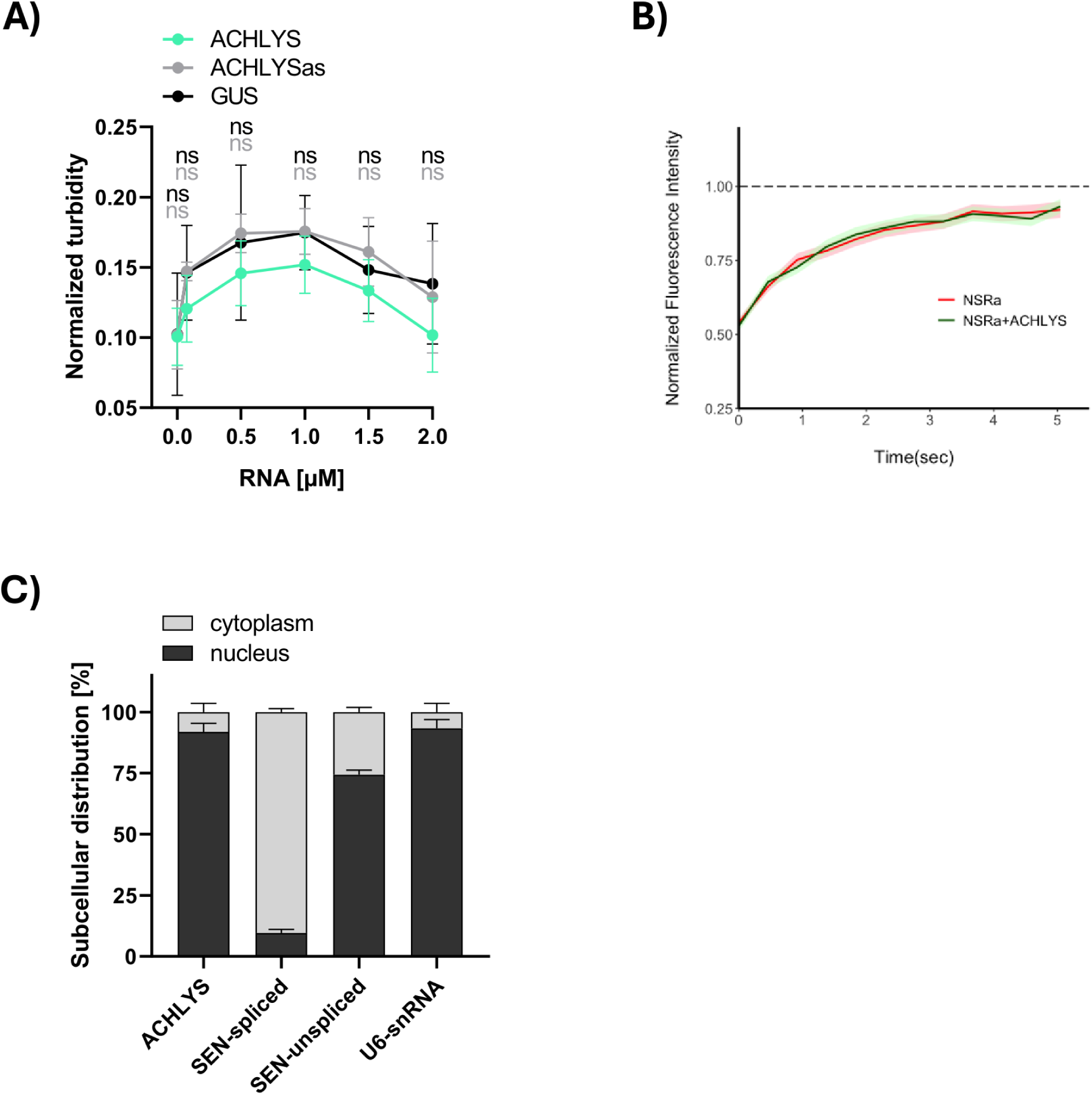
*In vitro* and *in vivo* formation of NSRa liquid liquid phase separation. **A)** *In vitro* phase separation of NSRa-RRM (6µM) in the presence of increasing *ACHLYS* (1234 nt), antisense (as) *ACHLYS* (1234 nt), APOLO, and GUS fragment (1247 nt). Mean and 95% confidence interval (n = 5-15, multiple unpaired, two-tailed T-test, ns = p > 0.05). **B)** Fluorescence Recovery After Photobleaching (FRAP) quantification of nsra pNSRa:NSRa-GFP and nsra pNSRa:NSRa-GFP OE-*ACHLYS* plants. Average recovery curve displays normalized fluorescence intensity in seconds after photo bleaching (n = 44-55). Fluorescence intensity is expressed relative to pre-photobleaching fluorescence. The shaded areas in color that accompany each curve represent the standard error for the values of each group. **C)** Subcellular localization of *ACHLYS* in OE-*ACHLYS* seedlings estimated by qPCR after fractionating cytoplasm and nucleus. SEN-spliced mRNA was used as a cytoplasmic RNA control. As a positive control for nuclear RNA localization, we utilized SEN-unspliced and U6 snRNA. The subcellular distribution was calculated as the percentage of 2^ΔCt-cytoplasm^+ 2 ^ΔCt-nucleus^. In OE-*ACHLYS* lines, *ACHLYS* accumulates in the nuclear fraction.

## Supplementary tables

**Table S1:** Gene ontology analysis of KD-*ACHLYS* DEG genes with log fold change -0.8 ≤ or 0.8 ≥.

| Pathways | Fold Enrichment | FDR | Genes |
| --- | --- | --- | --- |
| Response to hypoxia | 10.5 (6/267) | $1.2 \times 10^{-3}$ | RAV2, AT1G74940, ERF4, SZF1, WRKY70, NHL3 |
| Response to ethylene | 10 (6/282) | $1.4 \times 10^{-3}$ | RAV2, ORA47, ERF4, WRKY70, CRF1, GAMMA-VPE |
| Phosphorelay signal transduction system | 9.2 (5/254) | $6.6 \times 10^{-3}$ | RAV2, ORA47, ERF4, WRKY70, CRF1 |
| Response to abscisic acid | 5.6 (7/590) | $7.4 \times 10^{-3}$ | ESL1, COR413-PM1, KIN1, KIN2, BET9, ERF4, AT5G24240 |
| Response to hormone | 4.3 (15/1639) | $4.1 \times 10^{-4}$ | MYB77, RAV2, GASA6, ORA47, ERF4, WRKY70, CRF1, ESL1, COR413-PM1, ACS6, GAMMA-VPE, KIN1, KIN2, BET9, AT5G24240 |

**Table S2:**
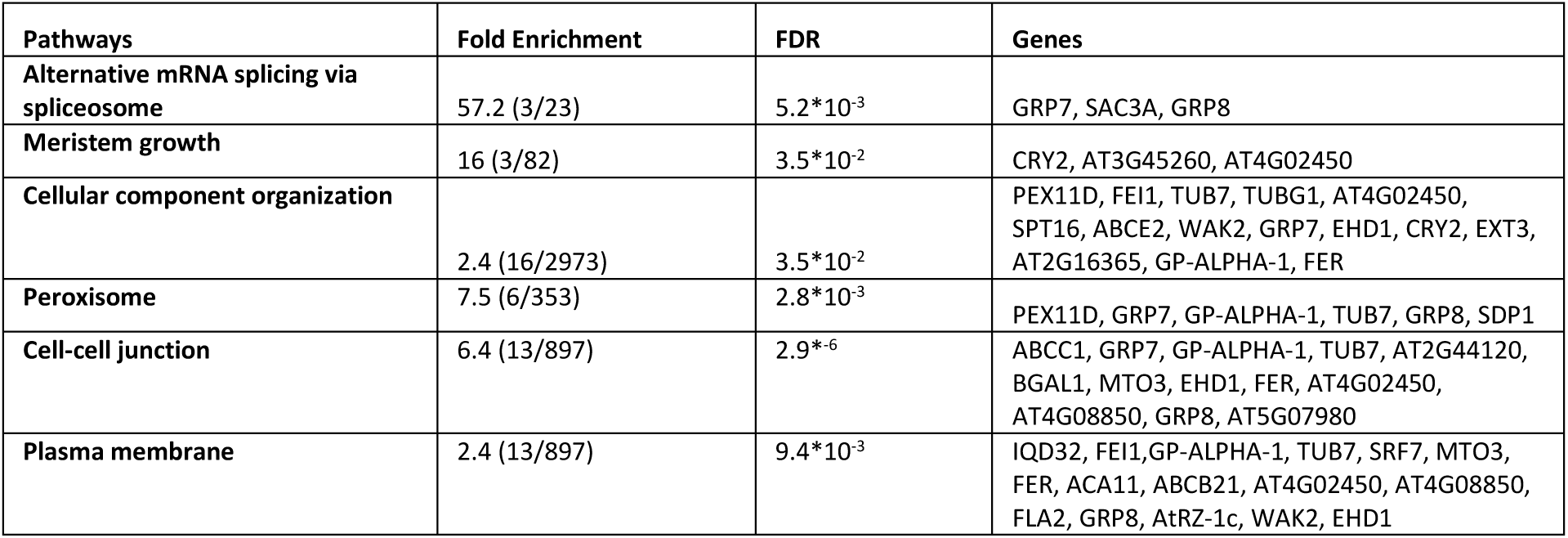
Gene ontology analysis of DAS events common in KD- and OE-*ACHLYS* with dPSI larger than 0.2 or smaller than -0.2 at least in one of the lines.

**Table S3:** List of plant lines utilized in this study.

| Name | Description | Reference |
| --- | --- | --- |
| ***ACHLYS-1*** | T-DNA insertion line SALK_083854, NASC order: N583854 | This study |
| ***ACHLYS-2*** | T-DNA insertion line SALK_043523, NASC order: N543523 |  |
| ***KD-ACHLYS-1*** | 35S:RNAi-*ACHLYS*/AT2G15292-1 |  |
| ***KD-ACHLYS-2*** | 35S:RNAi-*ACHLYS*/AT2G15292-2 |  |
| ***OE-ACHLYS-1*** | 35S:*ACHLYS*/AT2G15292-1 |  |
| ***OE-ACHLYS-2*** | 35S:*ACHLYS*/AT2G15292-2 |  |
| ***flail1*** | SMA3677, CRISPR mutant with a 229 bp deletion in the genomic region of *FLAIL* | Jin et al. 2023 |
| ***flail2*** | SMA3678, CRISPR mutant with a 343 bp deletion in the genomic region of *FLAIL* |  |
| ***flail3*** | SMA2648, T-DNA insertion line SAIL_645_C03, NASC order: N827819 |  |
| ***flail3* OE-GUS;** | SMA4477, *flail3* 35S:GUS |  |
| ***flail3* pFLAIL:FLAIL-1** | SMA4462, *flail3* pFLAIL:gFLAIL18 |  |
| ***flail3* pFLAIL:FLAIL-2** | SMA4464, *flail3* pFLAIL:gFLAIL88 |  |
| ***nsra*** | T-DNA insertion line SALK_003214 NASC order: N503214 | Bardou et al. 2014 |
| ***nsra* NSRa-GFP** | *nsra* complemented with pNSRa:NSRa-GFP |  |
| ***nsra* NSRa-GFP OE-*ACHLYS-1*** | *nsra* complemented with pNSRa:NSRa-GFP and 35S:*ACHLYS*/AT2G15292-1 | This study |
| ***nsra* NSRa-GFP OE-*ACHLYS-2*** | *nsra* complemented with pNSRa:NSRa-GFP and 35S:*ACHLYS*/AT2G15292-2 |  |
| ***nsra* NSRa-GFP OE-GRP7-2** | *nsra* complemented with pNSRa:NSRa-GFP and 35S:2-*GRP7*/AT1G04107 |  |

**Table S4:**
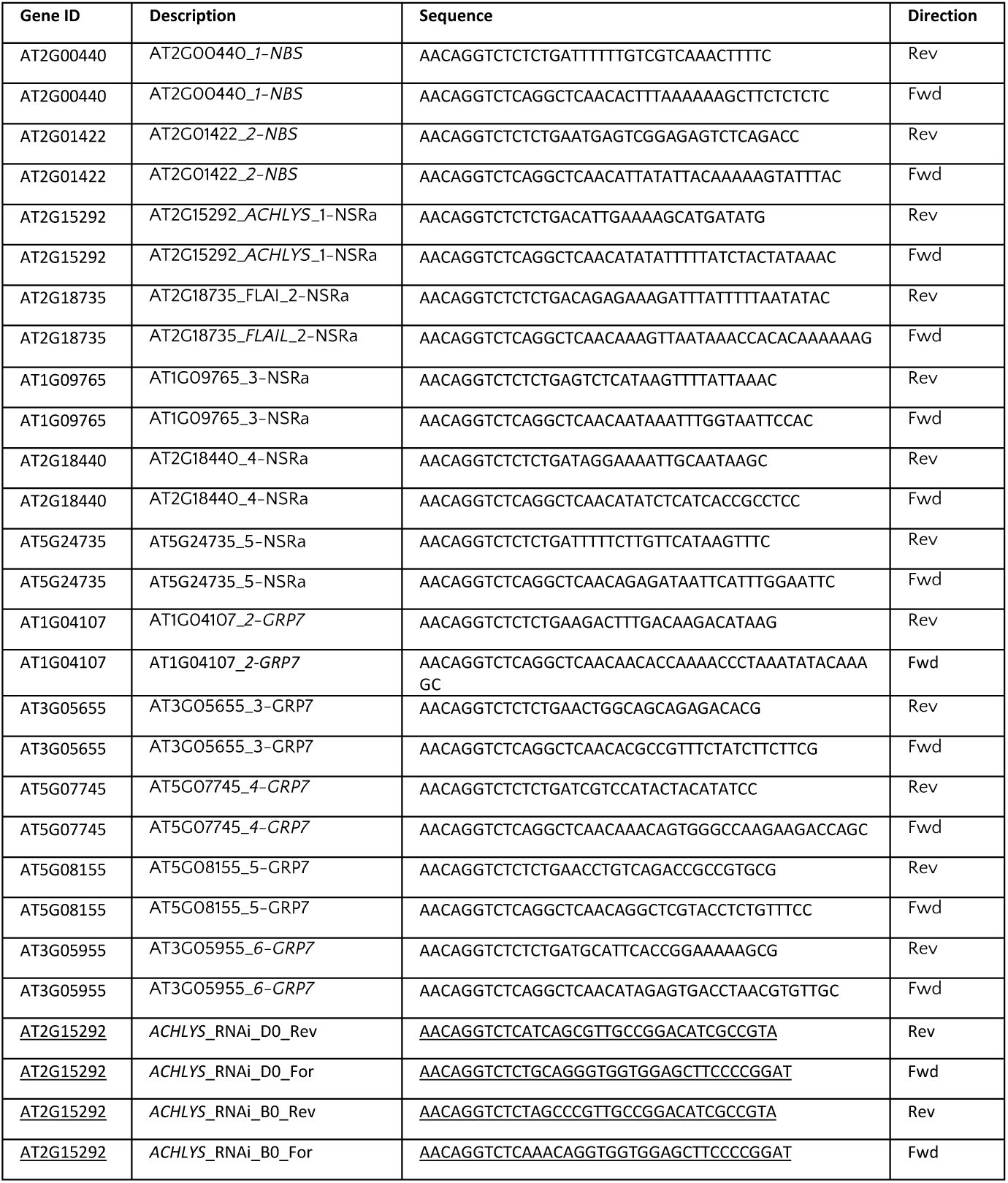
Primers used for cloning.

**Table S5:**
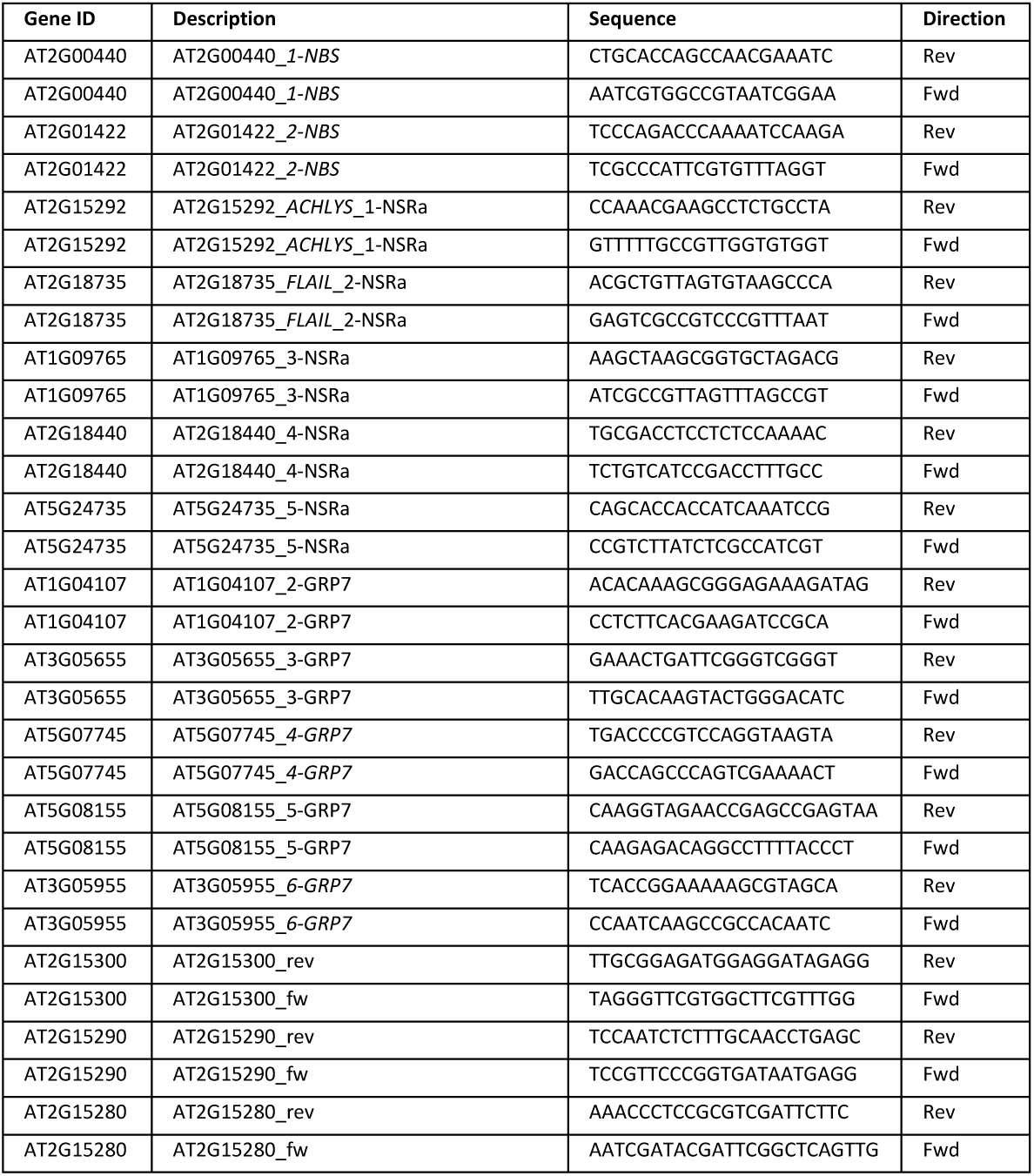
qPCR primers do determine gene expression levels.

**Table S6:**
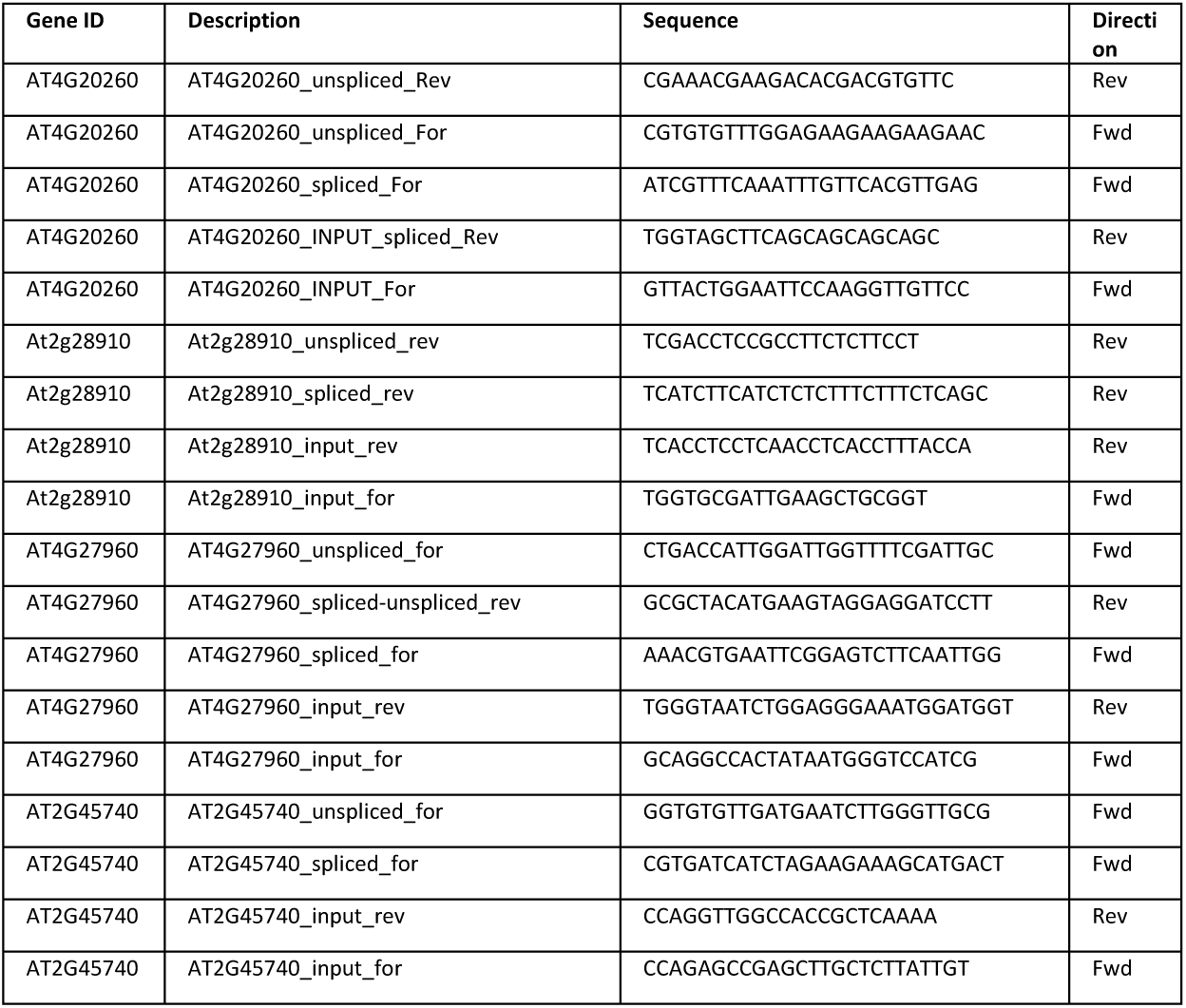
Splicing qPCR primers after transient lncRNA expression.

**Table S7:**
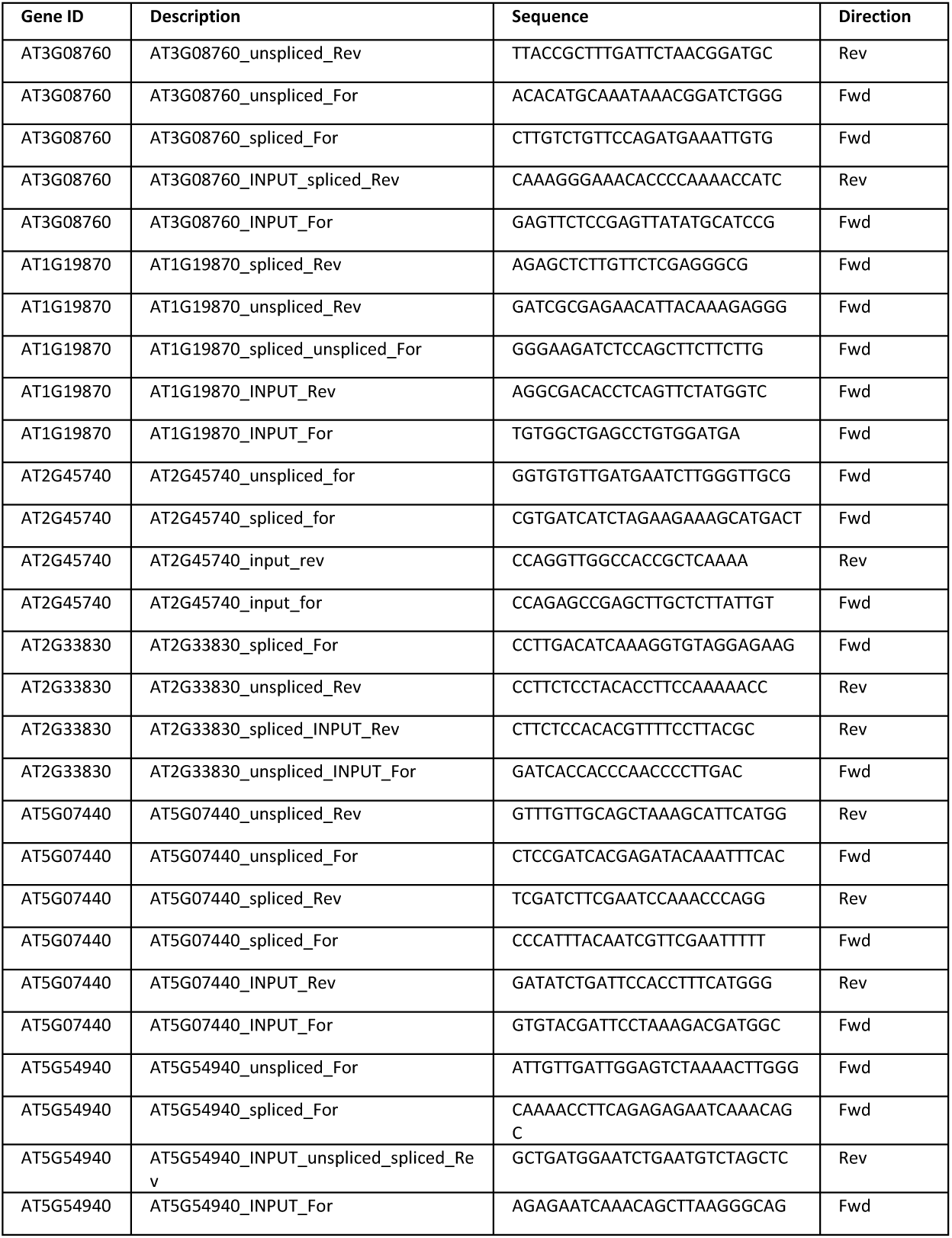
Splicing qPCR primers for KD- and OE-*ACHLYS*.

**Table S8:**
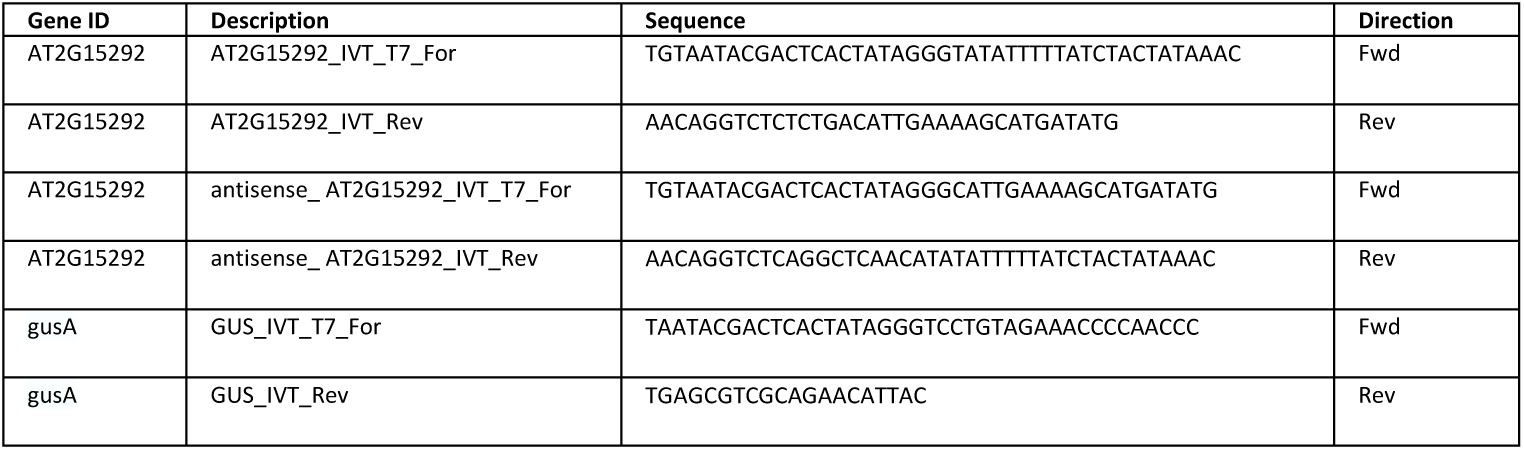
Primers for RNA *in vitro* transcription.

